# Spatial transcriptome sequencing of FFPE tissues at cellular level

**DOI:** 10.1101/2020.10.13.338475

**Authors:** Yang Liu, Archibald Enninful, Yanxiang Deng, Rong Fan

## Abstract

Formalin-fixed paraffin-embedded (FFPE) tissues are the most abundant archivable specimens in clinical tissue banks, but unfortunately incompatible with single-cell level whole transcriptome sequencing due to RNA degradation in storage and RNA damage in extraction. We developed an in-tissue barcoding approach namely DBiT-seq for spatially revolved whole transcriptome sequencing at cellular level, which required no tissue dissociation or RNA exaction, thus potentially more suited for FFPE samples. Herein, we demonstrated spatial transcriptome sequencing of embryonic and adult mouse FFPE tissue sections at cellular level (25μm pixel size) with high coverage (>1,000 genes per pixel). Spatial transcriptome of an E10.5 mouse embryo identified all major anatomical features in the brain and abdominal region. Integration with singlecell RNA-seq data for cell type identification indicated that most tissue pixels were dominated by single-cell transcriptional phenotype. Spatial mapping of adult mouse aorta, atrium, and ventricle tissues identified the spatial distribution of a variety of cell types. Spatial transcriptome sequencing of FFPE samples at cellular level may provide enormous opportunities in a wide range of biomedical research. It may allow us to exploit the huge resource of clinical tissue specimens to study human disease mechanisms and discover tissue biomarkers or therapeutic targets.

---

Clinical tissue samples are often stored as formalin fixed paraffin embedded (FFPE) blocks at room temperature, representing the most abundant resource of archived human specimens. For clinical histopathology and diagnostic purpose, tissue morphology is best preserved in FFPE as compared to other tissue banking methods, especially after prolonged storage^1,2^. Consequently, a huge volume of clinical FFPE tissue samples are readily available worldwide in hospitals and research institutions, which is a valuable source exploitable for retrospective tissue profiling and human disease research^3^. However, during the sample preparation and storage, nucleic acids including mRNAs in FFPE tissue often lost integrity and became partially degraded or fragmented^4^. In order to perform whole transcriptome analysis of FFPE samples using, for example, RNA sequencing (RNA-seq)^5^, a harsh chemical process for tissue decrosslinking, digestion, and RNA extraction is required, which unfortunately resulted in significant RNA degradation, damage, and loss. The bulk tissue digestion process also resulted in the loss of spatial and cellular information needed to trace the cellular origin of mRNAs^6, 7^.

Despite recent breakthroughs in massively parallel single-cell RNA sequencing (scRNA-seq) that have transformed all major fields of biological and biomedical research^8–10^, FFPE samples are not yet amenable to single-cell transcriptome sequencing using current techniques. Spatial transcriptomics emerged to address the limitation of scRNA-seq by retaining the spatial information of gene expression in the tissue context essential for a true mechanistic understanding of tissue organization, development, and pathogenesis. All early attempts of spatial transcriptomics were based on single-molecule fluorescence *in situ* hybridization(smFISH) or image-based in situ sequencing^11–13^. In order to measure the expression of mRNAs at the transcriptome level, it requires repeated hybridization and imaging cycles using high-end advanced fluorescence microscopy, which is technically demanding, costly, and time consuming. Moreover, most of these methods do not analyze RNAs base-by-base but rely on predesigned probes to detect known sequences only. It is highly desirable to harness the power of Next Generation Sequencing (NGS) to realize unbiased genome-wide profiling of spatial gene expression with high throughput and low lost. A barcoded solid-phase RNA capture approach was developed for coarse resolution (~100μm) spatial transcriptomics using DNA spot microarray^14^, which was recently improved to cellular resolution (~10μm) using self-assembled DNA barcode beads in Slide-seq and HDST^15, 16^. However, these NGS-based spatial transcriptomics methods were fundamentally limited by the requirement to de-crosslink FFPE tissues for RNA extraction, making it difficult to realize high-coverage transcriptome sequencing at cellular level.

We have developed high-spatial-resolution spatial omics sequencing vis deterministic barcoding in tissue (DBiT-seq)^17^, which was distinct from other NGS-based spatial transcriptome techniques in that it required no de-crosslinking for mRNA release and yielded high-quality transcriptome data from paraformaldehyde(PFA)-fixed tissue sections. Extending it to high-coverage spatial transcriptome sequencing of FFPE tissues at cellular level would be another major leap. Herein, we demonstrated spatially resolved whole transcriptome mapping of mouse embryo (E10.5) FFPE tissue samples with 25μm pixel size and identified all major tissue types in the brain and abdominal region at the cellular level. Integration with scRNA-seq data allowed for identification of 40+ cell types and revealed that most tissue pixels were dominated by single-cell transcriptome. Applying it to adult mouse heart (atrium and ventricle) and aorta tissues demonstrated high-coverage (>1000 genes per pixel) spatial transcriptome and the detection of sparse cell types in the cardiovascular tissues. This work represents a major leap forward to unlock the enormous resource of clinical histology specimens for human disease research.

The main workflow for FFPE samples is shown in **Figure 1a**. The banked FFPE tissue block was first microtomed into sections of 5-7 μm in thickness and placed onto a poly-L-lysine-coated glass slide. If the FFPE tissue sections were not to be analyzed right away, they should be stored at −80 °C prior to use in order to reduce RNA oxidative degradation by air exposure. Next, deparaffinization was carried out with standard xylene wash. Afterwards, the tissue section was rehydrated and permeabilized by proteinase K, and then post-fixed again with formalin. The deparaffinized tissue section ready for DBiT-seq exhibited a darkened and higher-contrast tissue morphology (**Figure 1b**). Then, the 1^st^ PDMS chip with 50 parallel channels was attached onto the tissue slide and a set of DNA barcode A1-A50 oligos were prepared in the reverse transcription (RT) mix and flowed through the channels to perform in tissue RT to produce cDNAs in situ with barcode A incorporated at the 3, end. Afterwards, the 1^st^ PDMS chip was removed and replaced with a 2^nd^ PDMS chip containing another set of 50 microchannels perpendicular to the first set of microchannels. Ligation was then performed in each of the channel by flowing a set of barcode B1-B50 oligos plus a universal ligation linker, which was complementary to the half-linker sequence in barcode A and B oligos in order to join them together in proximity to form full barcodes A-B. Thus, the ligation would only occur at the intersection of two flows where both barcode A and barcode B were present. Afterwards, the tissue was imaged and digested to collect cDNA to perform the downstream procedure including template switch, PCR amplification, and tagmentation to prepare the NGS library for paired-end sequencing.

**Figure 1.**
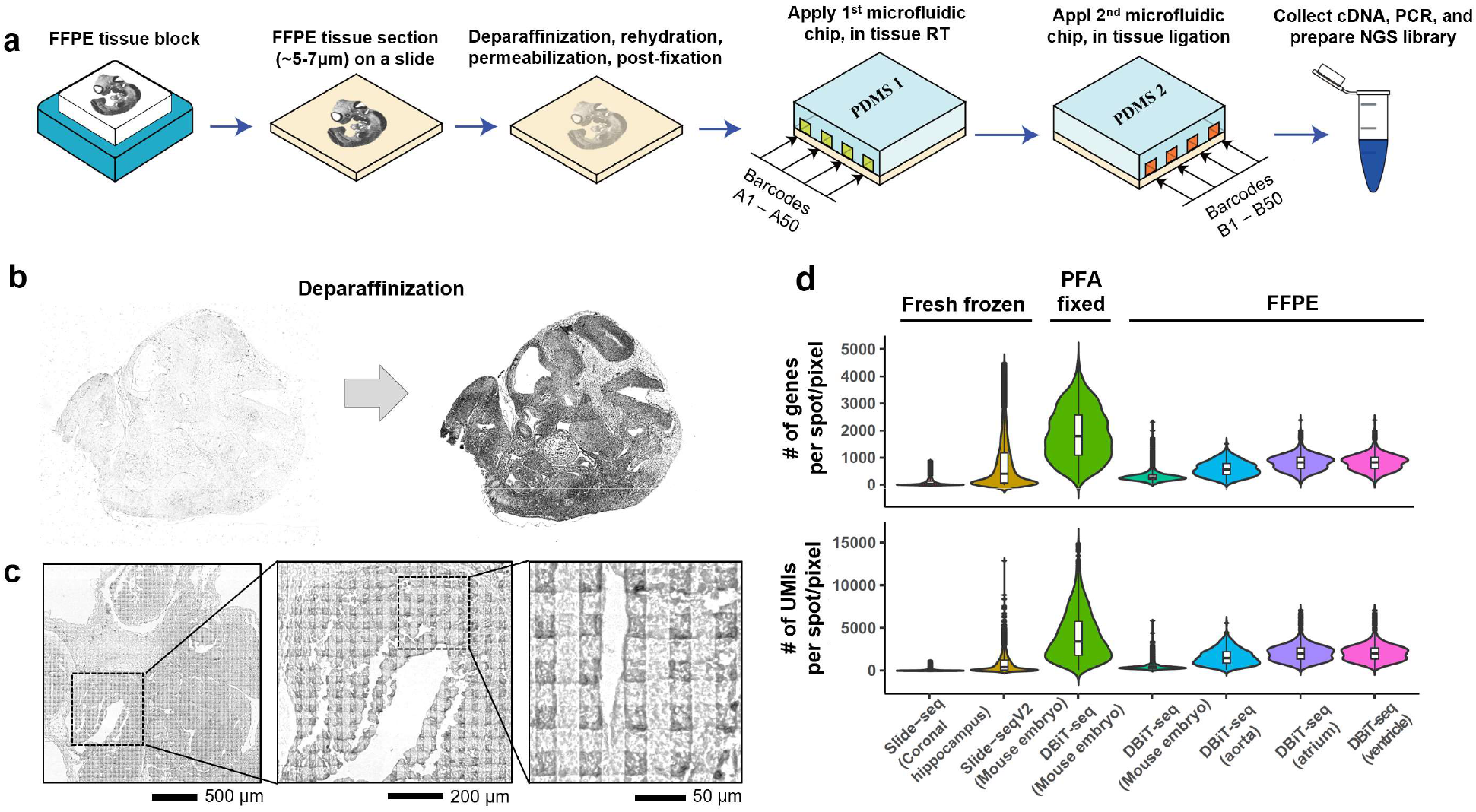
Workflow of DBiT-seq on FFPE samples. (a) Scheme of DBiT-seq on FFPE samples. FFPE tissue blocks stored at room temperature were sectioned into thickness of ~5-7 μm and placed onto a poly-L-lysine coated glass slide. Deparaffinization, rehydration, permeabilization and post-fixation were performed sequentially before placing the 1^st^ PDMS chip on the tissue section. Barcodes A1-A50 were flowed into the microchannels and the reverse transcription was carried out inside each channel. After washing, the 1^st^ PDMS was removed and a 2^nd^ PDMS chip with channels of perpendicular direction was attached on the tissue slide. Ligation reaction mix along with DNA Barcodes B1-B50 were pulled through each of the 50 channels by vacuum and incubated for 30 minutes to perform in situ ligation. Afterwards, the tissue section was digested with Proteinase K to collect cDNA for the downstream processes including template switch, PCR amplification, and tagmentation for NGS library preparation. (b) Deparaffinization of a E10 mouse embryo tissue. It maintained the original morphology with higher contrast after deparaffinization and the fine tissue features were readily discernable. (c) Deformation of tissue section after two sequential microfluidic flows of DBiT-seq. (d) Comparison of gene and UMI counts of our DBiT-seq data from FFPE samples with those obtained from other methods including Slide-seq, Slide-seqV2 and previous DBiT-seq data from PFA-fixed mouse embryo samples.

The attachment of PDMS chip to the tissue section was secured with a clamp set, and the clamping force could cause the deformation of tissue under the microfluidic channel walls. Therefore, after the application of two PDMS microfluidic chips onto the same tissue section in orthogonal directions, the slight deformation of tissue surface gave rise to a 2D grid of square features (**Figure 1c**), which allowed for the precise identification of individual DBiT-seq pixels and the corresponding location and morphology. The quality of cDNAs was evaluated by electrophoretic size distribution and compared between an archived FFPE mouse embryo sample and an FPA-fixed fresh frozen sample (**Figure S1a&b**). We noticed that the FFPE sample cDNA fragment size peaked between 400 and 500 bps, significantly shorter than that of the PFA-fixed fresh frozen sample which had the main peaks over 1000 bps. The average size was calculated to be ~600 bps for FFPE and ~1,400 bps for the PFA-fixed fresh frozen sample. This difference was due in part to the formalin cross-linking of RNA and proteins causing reduced accessible RNA segment length and the degradation of RNA during storage. Next, we assessed the quality of spatial transcriptome sequencing data based on total number of genes or unique molecular identifiers (UMIs) per pixel (**Figure 1d**). For FFPE samples, we found the results were variable among different experiments and sample types. For the mouse embryo samples, we obtained on average of 520 UMIs and 355 genes per pixel. For the mouse aorta sample, the average number of UMIs or genes per pixel were 1,830 and 663, respectively. For the adult mouse heart FFPE samples, we detected 3,014 UMIs and 1040 genes for atrium and 2,140 UMIs and 832 genes for ventricle. In comparison, we revisited the dataset of a PFA-fixed fresh frozen mouse embryo sample analyzed by DBiT-seq, which showed an average of 4,688 UMIs and 2,100 genes with the same pixel size (25μm). In order to validate the gene expression profile, we performed correlation analysis of the pseudo-bulk DBiT-seq data between FFPE and PFA-fixed fresh frozen mouse embryo tissue samples. The Pearson correlation coefficient R was ~0.88 (**Figure S1c**), which demonstrated a good agreement between the two types of experiments despite the difference in mapped tissue regions. The performance was also compared to spatial transcriptome mapping data from Slide-seq^15^ and Slide-seqV2^18^, which were obtained using unfixed fresh frozen mouse brain or embryo tissue samples.

Using an E10.5 mouse embryo FFPE tissue (**Figure 2a**), we conducted DBiT-seq on two adjacent sections to analyze two anatomic areas – the brain region (FFPE-1) and the abdominal region (FFPE-2), respectively. Using the Seurat package, clustering analysis of spatial pixel transcriptomes combining DBiT-seq data from both samples revealed 10 distinct clusters (**Figure 2b**). Mapping the clusters back to the spatial location identified spatially distinct patterns that agreed with the anatomical annotation (**Figure 2c**). Cluster 0 mainly represents the muscle structure in embryo. Cluster 3 covers the central nerve system including neural tube, forebrain and related nervous tissues. Cluster 4 is specific for ganglions, which comprises the brain ganglions and the dorsal root ganglions (**Figure 2c** right). High spatial resolution allows us to observe individual bone segments in the spine (cluster 6). Liver is largely shown as cluster 7. Heart comprises two layers of pixels with cluster 8 for myocardium and cluster 10 for epicardium. Cluster 9 is scattered within the neural tube region, probably representing a specific subset of neurons. These results demonstrated that high-spatial-resolution DBiT-seq could resolve fine tissue structures close to the cellular level. We further conducted GO analysis (**Figure 2d**) for each cluster, and the GO pathways matched well the anatomical annotation. The top 10 differentially expressed genes (DEG) were shown in a heatmap (**Figure S2**). We also conducted similar clustering analysis with each tissue sample as a separate dataset and the results revealed similar spatial patterns (**Figure S3**). DEGs for each cluster can be analyzed and compared (**Figure S4**). For example, *Stmn2* and *Mapt2*, which encode microtubule associated proteins and are important for neuron development, were mainly expressed in forebrain and the neural tube. *Fabp7*, a gene encoding the brain fatty acid binding protein, was expressed mainly in the hindbrain. Myosin associated genes, *Myl2, Myh7* and *Myl3*, were highly enriched in heart. *Slc4a1*, a gene related to blood coagulation, was detected extensively in liver, where most coagulation factors were produced. *Copx*, a heme biosynthetic enzyme encoding gene, was also produced in liver. *Afp*, which encodes alpha-fetoprotein, one of the earliest proteins synthesized by the embryonic liver, was observed exclusively in an organ-specific manner.

**Figure 2.**
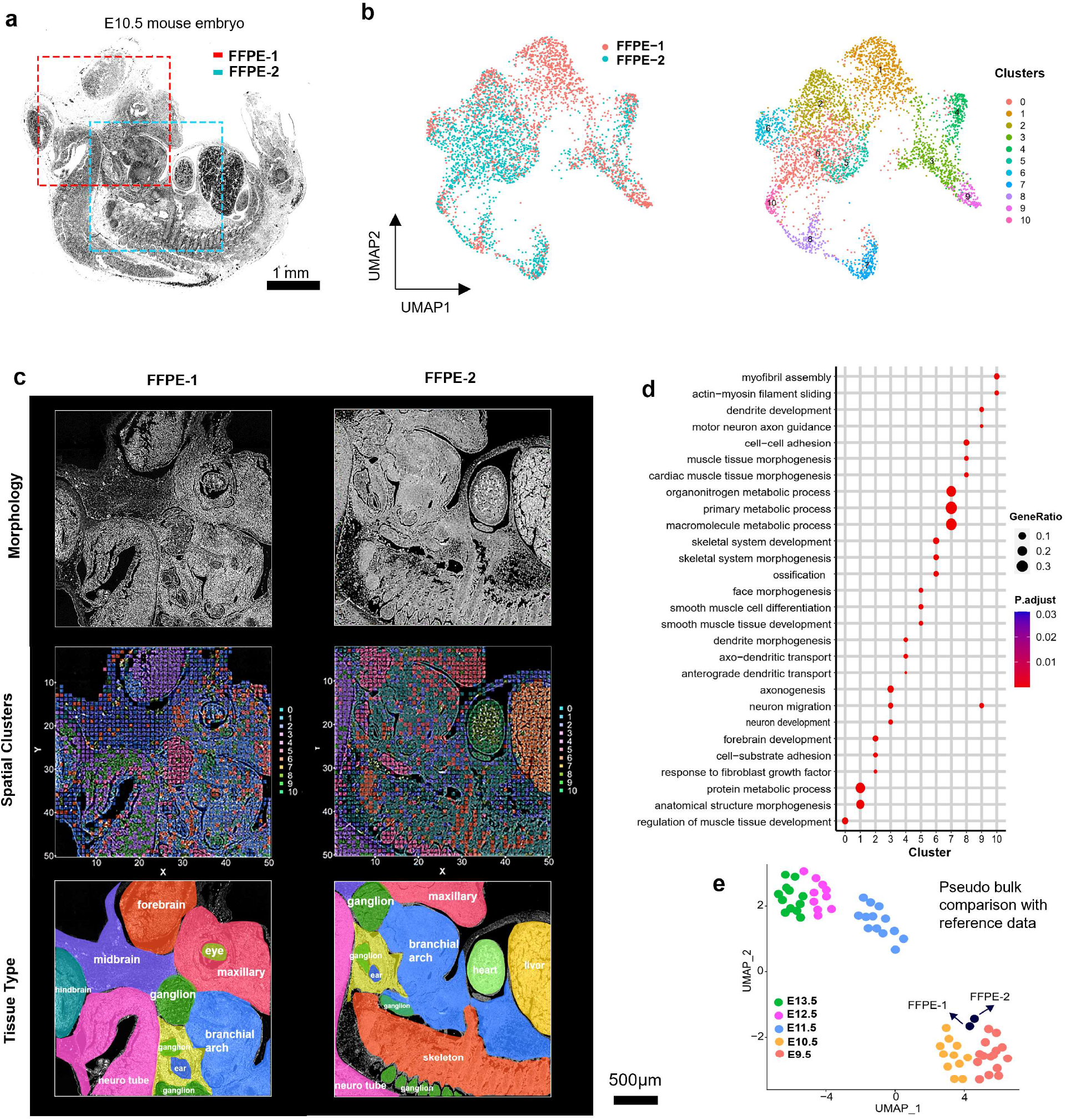
Spatial transcriptome analysis of FFPE tissue sections from an E10.5 mouse embryo. (a) Two tissue regions of FFPE mouse embryo were analyzed using DBiT-seq. One (FFPE-1) covered the brain region of the mouse embryo and the other (FFPE-2) covered the midbody abdominal region. Two separate tissue sections were used in this study. (b) UMAP visualization of combined pixels from FFPE-1 and FFPE-2 using Seurat package. Left: UMAP labelled by sample names; right: UMAP labelled by cluster numbers. Totally 11 clusters were identified. (c) Tissue morphology, anatomical annotation, and spatial mapping of the 11 clusters in (b). (d) GO enrichment analysis of all clusters (0-10). (e) Comparison of “pseudo bulk” data between DBiT-seq and scRNA-seq reference data. The aggregated transcriptome profiles of DBiT-seq from two FFPE samples conform well into scRNA-seq reference data from mouse embryos ranging from E9.5-E13.5(Cao et al., 2019).

We then applied SpatialDE, an unsupervised spatial differential gene expression analysis tool^19^, to the mouse embryo FFPE DBiT-seq data. It identified 30 spatial patterns for each of the two FFPE samples (**Figure S5&S6**). GO analysis of the SpatialDE-identified gene sets revealed the biological meaning of each spatial pattern. For example in FFPE-1 pattern 0 represents neural precursor cell proliferation and pattern 7 corresponds to eye morphogenesis. In FFPE-2, cluster 20 is specific for the heme metabolic process and cluster 26 strongly enriched in the heart tissue is for cardiac muscle contraction.

To identify the dominant cell type in each pixel, we performed integrated analysis of mouse embryo (E10.5) DBiT-seq data and scRNA-seq data from literature corresponding to the same developmental stage of mouse embryos^20^. We first compared the aggregated “pseudo bulk” data between DBiT-seq and scRNA-seq by unsupervised clustering (**Figure 2e**). In the UMAP plot, DBiT-seq data of FFPE-1 and FFPE-2 were in close proximity with the scRNA-seq data of E10.5 mouse embryo samples, which validated the FFPE DBiT-seq data for capturing the correct embryonic age even with lower coverage or the number of genes detected. We then performed the integrated analysis of these two types of data by combining the transcriptomes of all individual pixels from DBiT-seq with the transcriptomes of single cells for clustering analysis in Seurat after normalization with SCTransform^21^. The DBiT-seq pixels conformed to the clusters of scRNA-seq (**Figure 3a**), enabling the transfer of cell type annotations from single-cell transcriptomes to the spatial pixels and also to map different cell types back to spatial distribution (**Figure 3d**). In FFPE-1, cluster 3 mainly consisted of oligodendrocytes. Epithelial cells (cluster 4) and neural epithelial cells (cluster 13) were observed widely in epithelial glands. Interestingly, excitatory neurons (cluster 7) and inhibitory neurons (cluster 17) were both observed in the neural tube but forming a mixed pattern to fulfil their functionally distinct roles in transporting neurotransmitters. This integrative analysis answered an unresolved question in **Figure 2c** with regards to the specific subset of neurons observed in the neural tube by unsupervised clustering of DBiT-seq data alone. In FFPE-2, several organ-specific cell types were detected. For example, the primitive erythroid cells (cluster 14) crucial for early embryonic erythroid development and the transition from embryo to fetus in developing mammals were strongly enriched in liver^22^. Cardiac muscle cell types were observed mainly in the heart region in agreement with the anatomical annotation. In brief, integration with published scRNA-seq data could distinguish cell identity more robustly as compared to the differential gene expression and GO analysis of DBiT-seq data alone.

**Figure 3.**
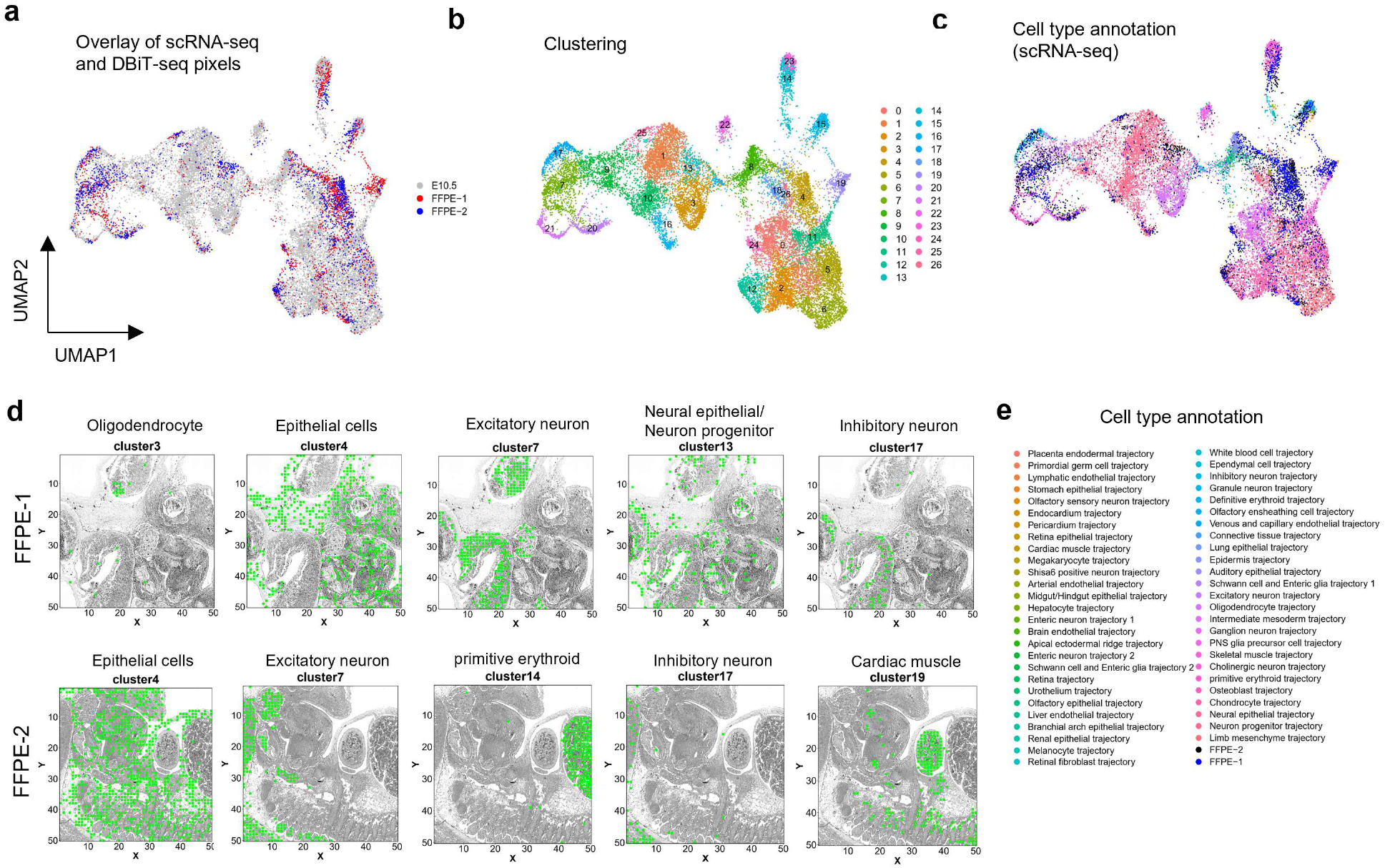
Integrative analysis of scRNA-seq and DBiT-seq (FFPE). (a) Integrative analysis of FFPE-1 and FFPE-2 DBiT-seq data with scRNA-seq data from mouse embryos ranging from E9.5-E13.5(Cao et al., 2019). The two samples conformed well in the scRNA-seq data. (b) UMAP of integrated data showing 26 distinct clusters. (c) UMAP clusters color coded for all cell types identified by scRNA-seq data combined with two FFPE sample spatial pixels. (d) Spatial expression of select clusters and corresponding cell types superposed on the tissue image. (e) List of all cell types identified in Figure 3c and spatial tissue pixels from two FFPE samples.

We next examined a mouse aorta FFPE tissue section (**Figure 4a**). The aorta tissue block was cross-sectioned, showing a thin wall of the artery along with the surrounding tissue. The heatmaps of gene and UMI counts (**Figure 4b**) showed more than 1,000 genes detected in 50% of the tissue pixels. Unsupervised clustering did not show distinct spatial patterns due to the lack of distinct tissue types and the dominance of specific cell types such as smooth muscle cells in this sample (**Figure S7b**). However, when integrated with scRNA-seq reference data from a mouse aorta^23^, we could identify six distinct cell types, including endothelial cells (ECs), arterial fibroblasts (Fibro), macrophages (Macro), monocytes(Mono), neurons and vascular smooth muscle cells (VSMCs). Most cells were ECs, VSMCs and Fibros. We also noticed that there was a layer of enriched smooth muscle cells in the artery wall, which were known to be the major cell type in a large artery^24^. We also performed automatic cell annotation using SingleR to analyze this aorta DBiT-seq data in comparison to the built-in reference database provided in the SingleR package based on scRNA-seq of mouse tissues (**Figure S7c**). It is worth pointing out that adipocytes that normally exist in the supporting tissue around the artery were readily identified. Meanwhile, the adipocyte-specific genes like *Adipoq* and *Aoc3* were observed to express at high levels in the surround tissue region (**Figure S7d**).

**Figure 4.**
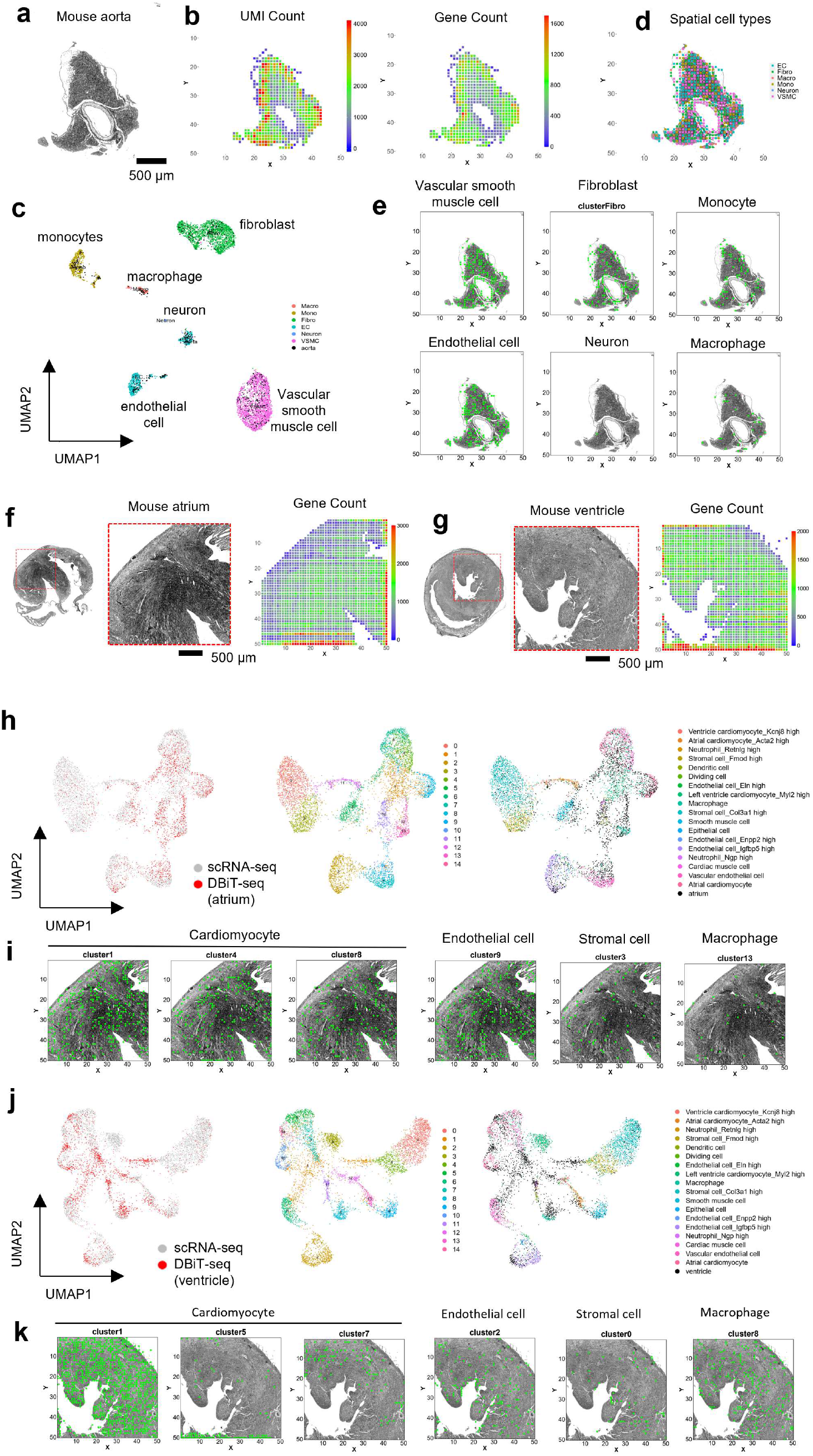
Spatial transcriptome mapping of adult mouse aorta, atrium and ventricle. (a) Bright field image of adult mouse aorta. Scale bar is 500 μm. (b) Spatial map of UMI and gene counts for each pixel. The average UMI count per pixel is ~1828 and gene count is ~664. (c) Clustering of spatial pixels with single-cell transcriptomes. The pixels conform to the clusters of scRNA-seq reference data. (d) Spatial distribution of major cell types annotated by integration with scRNA-seq data. These include endothelial cells (ECs), arterial fibroblasts (Fibro), macrophages (Macro), monocytes (Mono), Neurons and vascular smooth muscle cells (VSMCs). (e) Spatial distribution of individual cell types from Figure 4d. (f) Bright field image of a deparaffinized mouse atrium tissue section and the corresponding spatial gene count heatmap. (g) Bright field image of a deparaffinized mouse ventricle tissue section and the corresponding spatial gene count heatmap. (h) Clustering of the mouse atrium DBiT-seq data with scRNA-seq reference data. (i) Spatial distribution of representative annotated cells in atrium, see Figure S9 for the full panel. (j) Clustering of the mouse ventricle DBiT-seq data with scRNA-seq reference data. (k) Spatial distribution of representative annotated cells in ventricle, see Figure S9 for the full panel.

Lastly, we analyzed adult mouse atrium and ventricle FFPE samples using DBiT-seq (**Figure 4f&g**). Although cardiomyocytes only account for 30-40% of the total cell number in a heart, the volume fraction of cardiomyocytes can reach up to 70-80%^25^. Indeed, we observed the expression of muscle-related genes like *Myh6* extensively throughput the cardiac tissue (**Figure S8a**), which encodes a protein known as the cardiac alpha (α)-myosin heavy chain. The fact that there is a large volume of cardiomyocytes in this tissue posed a challenge for spatial expression pattern analysis due to the dominance of one specific cell type and the lack of distinct anatomic landmarks. Unsupervised clustering of the DBiT-seq pixels from atrium and ventricle using Seurat could not resolve highly distinct clusters (**Figure S8b&c**). However, when integrated with scRNA-seq reference data from the mouse hearts^26^, DBiT-seq pixels of atrium and ventricle conformed rather well to single-cell transcriptional clusters and revealed a total of 15 clusters (**Figure 4h&j**). These clusters were then annotated using the cell types defined by scRNA-seq (**Figure S9**). The results confirmed that cardiomyocytes were still the main cell type in this tissue and observed across multiple spatial clusters (**Figure 5d&f**), for example, clusters 1, 4, and 8 in the atrium. A significant number of endothelial cells were observed, presumably corresponding to coronary microvasculature in myocardium. Other cell types, like stromal fibroblasts and macrophages were observed presumably in the interstitial space of cardiomyocyte fibers in the mouse heart.

In summary, we demonstrated spatially resolved transcriptome sequencing of FFPE tissue sections with 25μm pixel size. The data quality in terms of the number of UMIs and genes detected was lower than that from PFA-fixed frozen sections, but still yielded highly meaningful results with ~1,000 genes per pixel achieved across whole transcriptome, which was comparable to other high-spatial-resolution (10 or 20μm spot size) spatial transcriptome technologies^15, 16^ that are currently compatible with fresh frozen samples only. Applying our technology to mouse embryo FFPE tissues resulted in the identification of 11 spatial patterns that agreed with anatomical annotations. Integration with published scRNA-seq data further improved cell type identification and revealed that most spatial tissue pixels were dominated by single-cell transcriptome. We further analyzed adult mouse aorta, atrium and ventricle FFPE tissue samples and revealed a wide range of cell types localized in the interstitial space of myocardium or the perivascular supporting tissue. As FFPE samples are widely available and represent the most abundant format of archivable clinical tissue samples, we envision that this work will open up new opportunities to revisit the huge resource of clinical tissue banks to study the mechanisms of pathophysiology and to discover new targets for diagnosis and treatment of human diseases.

## Acknowledgements

This research was supported by Packard Fellowship for Science and Engineering (to R.F.), Stand-Up-to-Cancer (SU2C) Convergence 2.0 Award (to R.F.), and Yale Stem Cell Center Chen Innovation Award (to R.F.), National Science Foundation CAREER Award CBET-1351443 (R.F.), National Institutes of Health grants U54 CA209992 (Sub-Project ID: 7297 to R.F.), R01 CA245313 (R.F.), R33 CA196411 (R.F.), R33 CA246711 (R.F), and UG3CA257393, to R.F.). Y.L. was supported by the Society for ImmunoTherapy of Cancer (SITC) Fellowship. The molds for making microfluidic chips were fabricated at the Becton Nanofabrication Center at the Yale University. We used the service provided by the Genomics Core of Yale Cooperative Center of Excellence in Hematology (U54DK106857). Next-generation sequencing was conducted at Yale Stem Cell Center Genomics Core Facility which was supported by the Connecticut Regenerative Medicine Research Fund and the Li Ka Shing Foundation. It was also conducted using the sequencing facility at the Yale Center for Genomic Analysis (YCGA).

## Author contributions

Conceptualization: Y.L, R.F.; Methodology, Y.L., A.E., and Y.D.; Experimental investigation, Y.L., A.E., and Y.D.; Data Analysis, Y.L., A.E., and Y.D. and R.F.; Writing – Original Draft, Y.L. and R.F.; Writing – Review and Editing, Y.L., A.E., Y.D. and R.F..

## Supplementary information

Supplementary Information can be found online at [to be inserted, SI is also provided as part of the manuscript submission].

## Methods

### Fabrication of microfluidic device

Soft lithography was used to produce the PDMS (polydimethylsiloxane) microfluidic device. The chrome mask was printed by the company Front Range Photomasks (Lake Havasu City, AZ) with high resolution (2 μm). Upon receiving, chrome mask was cleaned using acetone before use to remove any dirt or dust. The negative photoresist SU-8 (SU-8 2025) based mold was fabricated according to manufacturer’s (MicroChem) recommendations using a precleaned silicon wafer substrate. The final SU-8 layer of the mold had a thickness of ~25 μm and a channel width of 25 μm. The fabrication of PDMS microfluidic chips were through a replication molding process. The GE RTV PDMS part A and part B were mixed thoroughly with a 10:1 ratio and poured onto the mold. After degassing for 30min, the PDMS was cured in a 75 °C oven for 2 hours. The cured PDMS slab was then cut and punched with inlet and outlet holes using a 2 mm diameter puncher. The acrylic clamps (rectangle, 22 mm × 40 mm) to strength the attachment of PDMS to glass slide were fabricated using a laser cutter.

### Tissue Handling

FFPE samples of adult mouse and mouse embryo were obtained from Zyagen (San Diego, CA). According to Zyagen protocol, the mice used in this project were purchased from Charles River Laboratories. Adult mice were sacrificed upon arrival, and the aorta, atrium and ventricle were collected. The embryo (E10.5) were collected the day the pregnant mouse was received. The FFPE tissues were processed following standard protocols, which includes fixation (10% formalin), dehydration (ethanol series: 70%-100%), clearing (100% xylene), paraffin infiltration and embedding. FFPE sample was sectioned with a thickness of 5-7 μm and placed onto a poly-L-lysine coated glass slide. After receiving the sectioned FFPE slides, the tissue sections were stored at −80 °C in a sealed bag until use.

### Deparaffinization of tissue section

Prior to deparaffinization, adult mouse or mouse embryo FFPE tissue slides were first baked at 60°C for 1 hour to ensure that the tissue sections were properly attached to glass slide. Deparaffinization was performed by two times washing with Xylene (100%) for 5 minutes each. To remove the remaining xylene, the section was washed 5 minutes with 100% ethanol. Tissue was then rehydrated by immersing in 90%, 70% and 50% ethanol for 5 minutes each, and finally placed in PBS with 0.1% Tween-20. The tissue was permeabilized for 5 minutes with Proteinase K 7.5μg/ml in PBST and fixed in 4% formaldehyde with 0.2% Glutaraldehyde for 20 minutes.

### DNA oligo design

Two sets of DNA barcodes (A1:A50 and B1:B50) were used in this study. Barcode A1:A50 had three different functional regions: a 16-mer poly-T region, an 8-mer spatial barcode region (mark Y-axis location) and a 15-mer ligation linker region (See example Barcode A below). Barcode A1:A50 served as the RT primer and was loaded into each of the 50 channels of the 1^st^ PDMS along with reverse transcription mix. The resulting cDNA products were then ligated to the barcode B1:B50 during the ligation process. There were four different functional parts in barcode B: a 15-mer ligation linker, an 8-mer spatial barcode region(mark X-axis location), a 10-mer unique molecular identifier (UMI), and a PCR handle functionalized with biotin, which is used for purification purpose. Before loading into the 2^nd^ PDMS, barcode B was first annealed to a complementary ligation linker strand and then mixed with DNA ligase reaction mix. The ligation product baring the x and y location information was then extracted and processed with downstream steps. There were theoretically 2,500 pixels in a tissue region of 2.5 mm × 2.5 mm square.

DNA barcode Examples

Barcode A1:

/5Phos/AGGCCAGAGCATTCGAACGTGATTTTTTTTTTTTTTTTVN

Barcode B1: /5Biosg/CAAGCGTTGGCTTCTCGCATCTNNNNNNNNNNAACAACCAATCCACGTGCTTGAG

DNA Oligos and barcodes used in this paper were listed in Table S1 and all other reagents were listed as Table S2 and S3.

### In tissue reverse transcription with Barcode A

The deparaffinized tissue section was blocked by 1% BSA solution in PBS plus RNase inhibitor (0.05U/μL, Enzymatics) for 30 minutes at room temperature. After 3-times washing with 1X PBS and 1-time wash with water, the 1^st^ PDMS slab with 50 channels was placed on the glass slide, covering the interested tissue region. The brightfield image (10x, Thermo Fisher EVOS fl microscope) was recorded and used later for the identification of pixel locations. Afterwards, an acrylic clamp with screws was clamped against the center tissue region of interest.

The Reverse Transcription solution (225 μL) was first prepared by mixing:

50 μL of RT buffer (5X, Maxima H Minus kit),
32.8 μL of RNase free water,
1.6 μL of RNase Inhibitor (Enzymatics),
3.1 μL of SuperaseIn RNase Inhibitor (Ambion),
12.5 μL of dNTPs (10 mM, Thermo Fisher),
25 μL of Reverse Transcriptase (Thermo Fisher),
100 μL of 0.5X PBS with Inhibitor (0.05U/μL, Enzymatics).

After vortex mixing, the RT solution was aliquoted into 50 different tubes, with each tube a 4.5 μL solution. Then, into each tube, a 0.5 μL of barcodes A (A1-A50) (25 μM) was added and mixed thoroughly. The 50 tubes of 5 μL of RT reaction solution were loaded into the 50 inlets (each can hold >10 μL solution) on the PDMS. In order to fill up the channel and remove air bubbles, the solution was pulled through each of the 50 channels with vacuum continuously for 3 minutes. The chip was then put into a wet box and incubated at room temperature for 30 minutes and then at 42 °C for another 1.5 hours. After RT, the channels were cleaned up by 1X NEB buffer 3.1(New England Biolabs) with 1% RNase inhibitor (Enzymatics) continuously for 10 minutes. Finally, the clamp and PDMS were removed from the tissue slide. The slide was quickly dipped in water and dried with air.

### In tissue ligation with Barcode B

The 2^nd^ PDMS slab with channels perpendicular to the 1^st^ PDMS was attached to the dried slide with care. A brightfield image was taken (10x, Thermo Fisher EVOS fl microscope) and the same clamp was used here to press the PDMS against the tissue. We then prepared 115.8 μL ligation mix by adding the following reagents into a 1.5 mL Eppendorf tube.

69.5 μL of RNase free water,
27 μL of T4 DNA ligase buffer (10X, New England Biolabs),
11 μL T4 DNA ligase (400 U/μL, New England Biolabs),
2.2 μL RNase inhibitor (40 U/μL, Enzymatics),
0.7 μL SuperaseIn RNase Inhibitor (20 U/μL, Ambion),
5.4 μL of Triton X-100 (5%).

DNA barcode B was first annealed with ligation linker by adding 25 μL of Barcode B (100 μM), 25 μL of ligation linker (100 μM) and 50 μL of annealing buffer (10 mM Tris, pH 7.5 – 8.0, 50 mM NaCl,1 mM EDTA). 5 μL ligation reaction solution (totally 50 tubes) was prepared by adding 2 μL of ligation mix, 2 μL of NEB buffer 3.1(1X, New England Biolabs) and 1 μL of each DNA barcode B and ligation linker mix (B1-B50, 25 μM) and then loaded into each of the 50 channels with vacuum. The chip was kept in a wet box and incubated at 37 °C for 30 minutes. After washing by flowing 1X PBS with 0.1% Triton X-100 and 0.25% SUPERase In RNase Inhibitor for 10 minutes, the clamp and PDMS were removed, and the dried slide was ready for tissue digestion.

### Tissue digestion

After removing the 2^nd^ PDMS, the tissue section was dipped in water and dried with air before taking the final brightfield image. Afterwards, we prepared proteinase K lysis solution, which contains 2 mg/mL proteinase K (Thermo Fisher), 10 mM Tris (pH = 8.0), 200 mM NaCl, 50 mM EDTA and 2% SDS. We then covered the tissue region of interest with a square well PDMS gasket and then loaded around ~25 μL of lysis solution into it. The lysis was performed at 55 °C for 2 hours in a wet box. The tissue lysate was collected into a 1.5 mL Eppendorf tube and purified using streptavidin beads (Dynabeads MyOne Streptavidin C1 beads, Thermo Fisher) or stored at −80 °C until use.

### cDNA extraction

Before extraction, RNase free water was first added into the lysate to bring the total volume up to 100 μL. 5 μL of PMSF (100 μM, Sigma) was added to the lysate and incubated for 10 minutes at room temperature to inhibit the activity of Proteinase K. Meanwhile, the magnetic beads were cleaned three times with 1X B&W buffer with 0.05% Tween-20 and dispersed into 100 μL of 2X B&W buffer (with 2 μL of SUPERase In Rnase Inhibitor). After adding 100 μL of the cleaned streptavidin beads suspension to the lysate, the mixture was incubated for 60 minutes at room temperature with gentle shaking. Afterwards, the beads were washed twice with 1X B&W buffer and 1X Tris buffer (with 0.1% Tween-20) once.

### Template switch

After cleaning, the beads were resuspended into 132 μL of the template switch reaction mix, which consists of:

44 μL 5X Maxima RT buffer (Thermo Fisher),
44 μL of 20% Ficoll PM-400 solution (Sigma),
22 μL of 10 mM dNTPs each (Thermo Fisher),
5.5 μL of RNase Inhibitor (Enzymatics),
11 μL of Maxima H Minus Reverse Transcriptase (Thermo Fisher),
5.5 μL of a template switch primer (100 μM).

Template switch was performed first at room temperature for 30 minutes and then at 42 °C for 90 minutes. After reaction, the beads were pulled down using the magnetic stand and rinsed once with 500 μL 10 mM Tris plus 0.1% Tween-20, and then cleaned with 500 μL RNase free water.

### PCR amplification

There are two separate PCR processes. In the first PCR, the cleaned beads with template switched cDNAs were first resuspended into the PCR mix, which contains 110 μL Kapa HiFi HotStart Master Mix (Kapa Biosystems), 8.8 μL of 10 μM stocks of primers 1 and 2, and 92.4 μL of water. The mix were aliquoted into 4 different PCR tubes, with each ~50 μL of solution. Then, PCR reaction was performed with the following steps: incubate at 95°C for 3 mins, then cycle five times at

98°C for 20 seconds,
65°C for 45 seconds,
72°C for 3 minutes.

After reaction, the beads were removed, and the supernatant was collected and pipetted into 4 new PCR tubes. A second PCR was performed by first incubating at 95°C for 3 minutes, then cycled 20 times at

98°C for 20 seconds,
65°C for 20 seconds,
72°C for 3 minutes.

The PCR product was kept in 4°C until next step.

### Sequencing library preparation

To remove remaining PCR primers, the PCR product was purified using the Ampure XP beads (Beckman Coulter) at 0.6x ratio following standard protocol. The purified cDNA was then quantified by an Agilent Bioanalyzer High Sensitivity Chip. A Nextera XT Library Prep Kit (Illumina, FC-131-1024) was used to prepare the sequencing library. 1 ng of the cDNA was used as the starting material, and the library preparation is following manufacture protocols. The library was then analyzed by bioanalyzer again and sequenced using a HiSeq 4000 sequencer with pair-end 100×100 mode.

### DBiT-seq data pre-processing

Read 1 of the raw sequencing data contains the transcriptome information, while Read 2 holds the UMI, Barcode A and Barcode B. Following ST pipeline v1.7.2^27^, Read 1 was trimmed, filtered, STAR mapped (STAR version 2.6.0a) against the mouse genome(GRCh38) and annotated using Gencode release M11. Most of the default parameters were used when running ST pipeline, except that the “--min-length-qual-trimming” was set to 10. The final expression matrix has the location info as rows and gene expression levels as columns. R package “ggolot2” was used to plot the spatial heatmaps for pan-mRNA or individual genes.

### UMI and Gene Counts comparison with other techniques

To compare with other NGs based spatial RNA-seq technique (Fresh Frozen tissue sections), we downloaded published data:

10X Visium: 151673_filtered_feature_bc_matrix.h5 (human cortex)
Slide-seq: Puck_180413_7(coronal hippocampus)^15^
Slide-seqV2: Puck_190926_03 (mouse embryo)^18^
DBiT-seq: Mouse embryo Brain E11 25μm resolution^17^

The total UMI and Gene counts were calculated for each of the spots(pixels) and then the violin plots for each technique were plotted side-by-side.

### Pseudo bulk comparison with reference

The pseudo bulk data for FFPE samples were obtained by summing counts for each gene in each sample and divided by the sum of total UMI counts, and further multiplied by 1 million. Similarly, the pseudo bulk data was calculated using the E9.5-E13.5 embryo scRNA-seq data from Reference paper^20^.

### Clustering with Seurat

We used Seurat V3.2^28, 29^ to analyze the spatial transcriptome data of all the FFPE samples. Data integration and normalization were performed with the SCTransform workflow. The top 3,000 variable features were selected when doing data integration. For PCA analysis and UMAP visualization, the dimensions were set to 10, and the clustering resolution was set to 0.8. Differentially expressed genes for each cluster was obtained by comparison of cells in individual clusters against all remaining cells.

### SpatialDE analysis

To study spatial patterns of gene expression, SpatialDE, an unsupervised automatic expression analysis tool was conducted for both adult heart and mouse embryo samples. Following standard workflow, SpatialDE identified >15 distinct spatial patterns in the mouse embryo sample. The results agree with Seurat pixel-based clustering results.

### Cell type annotation

Cell type annotation was achieved by integration analysis (Seurat V3.2, SCTransform) combining the spatial transcriptome data of FFPE samples and the corresponding published scRNA-seq reference. After clustering, the spatial pixel data conformed well with the scRNA-seq data, and thus the cell types were assigned based on the scRNA-seq cell type annotation for each cluster (if two cell types presented in one cluster, the major cell types were assigned). SingleR is also used for aorta sample annotation with the built-in reference “MouseRNAseqData”^30^.

### GO analysis

GO analysis was completed using the “GO Enrichment Analysis” module at http://geneontology.org/ with default settings. The biological process was ranked by the gene ratio and the top 3-5 biological process were plotted using the “dotplot” function in ggplot2.

### Data sharing and codes

Data is available at: https://www.ncbi.nlm.nih.gov/geo/query/acc.cgi?acc=GSE156862. Codes for data analysis are available (https://github.com/rongfan8/DBiT-seq_FFPE).

## SUPPLEMENTARY INFORMATION

### Supplementary Figures

**Figure S1.**
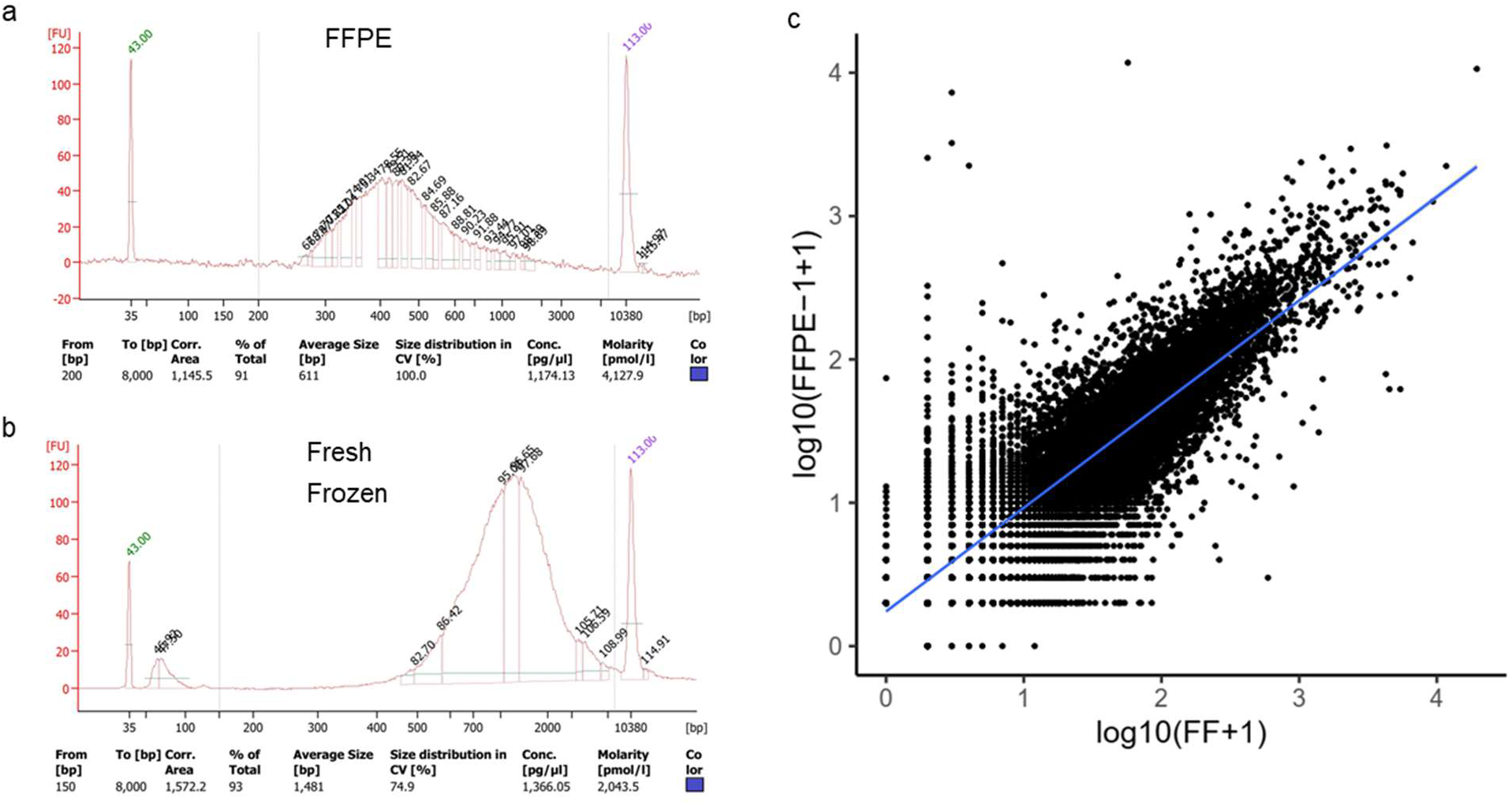
Size distribution of cDNAs from FFPE and fresh frozen mouse embryo samples and correlation of sequencing data between FFPE sample and PFA-fixed frozen samples. (a) FFPE cDNA bioanalyzer data. (b) Fresh Frozen cDNA bioanalyzer data. (c) Correlation of sequencing data between FFPE and fresh frozen samples. Pearson correlation coefficient R = 0.88.

**Figure S2.**
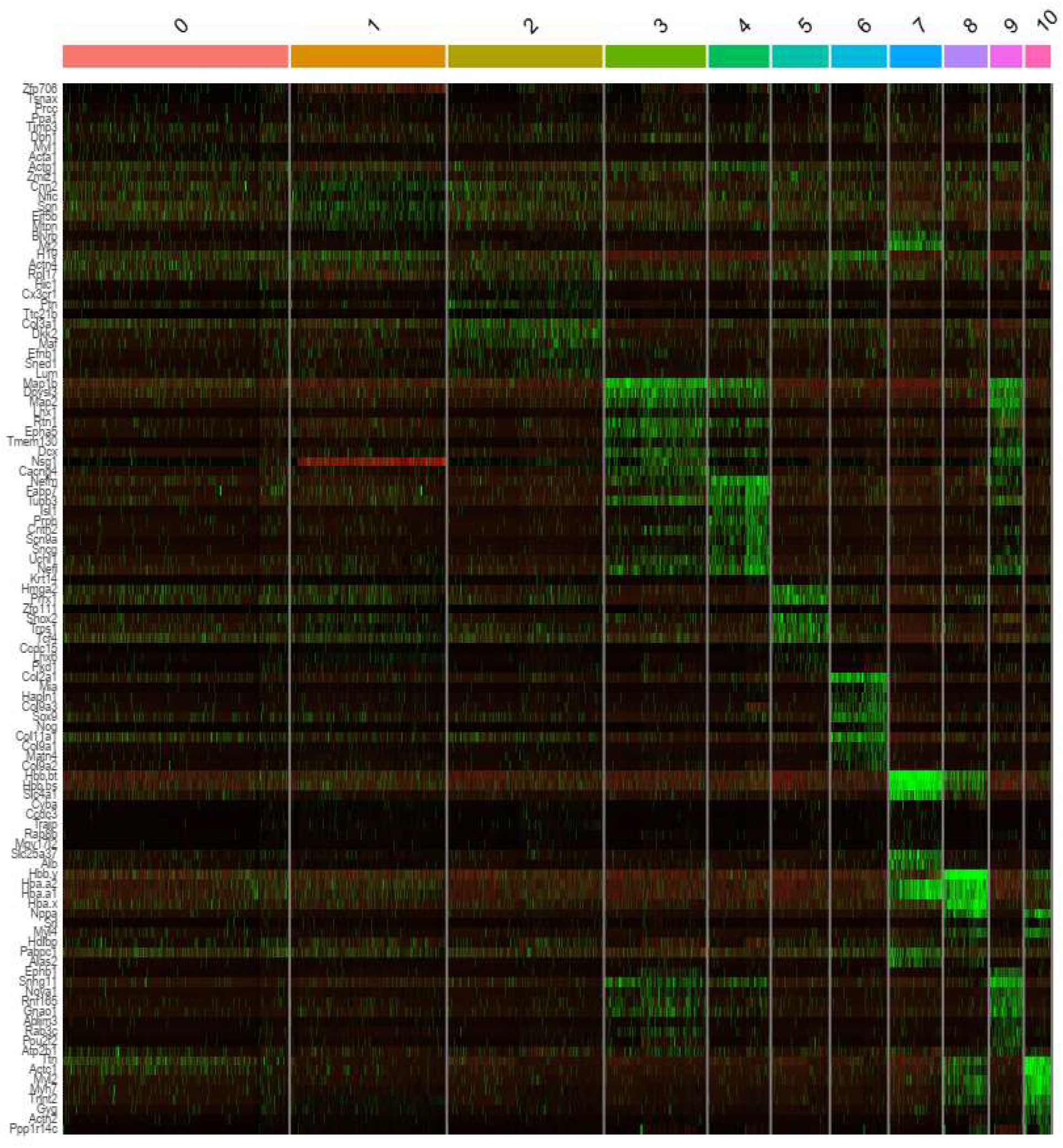
Differential Expressed Gene (DEG) heatmap of FFPE-1 and FFPE-2 (combined). This heatmap is a summary of top 10 DEGs in each cluster.

**Figure S3.**
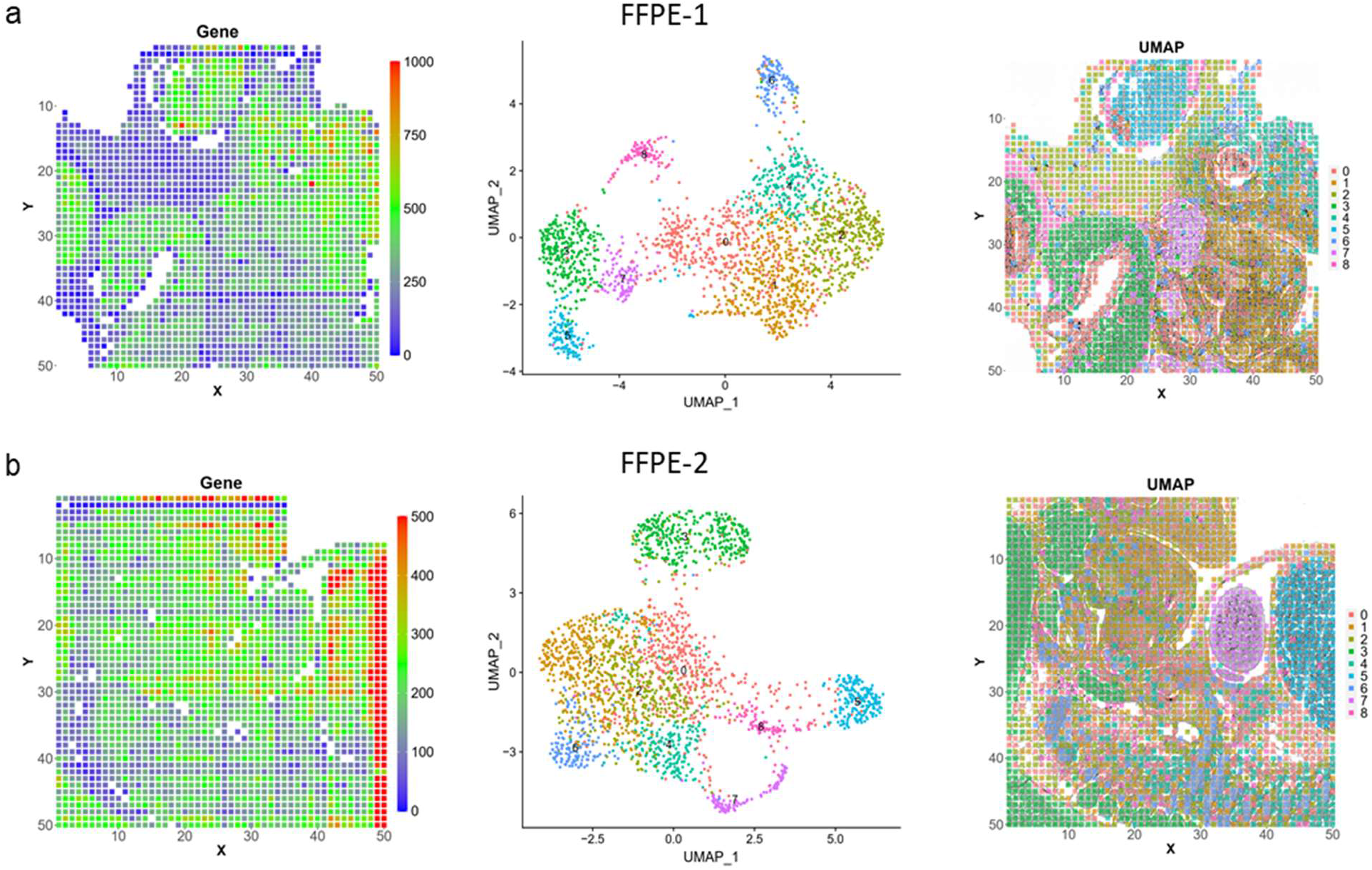
Clustering analysis of FFPE-1 and FFPE-2. (a) Clustering analysis of FFPE-1 only. Left: gene counts heat map; middle: UMAP of all FFPE-1 pixels; right: Spatial mapping of UMAP clusters. 8 clusters were observed. (b) Clustering analysis of FFPE-2. Left: gene counts heat map; middle: UMAP of all FFPE-1 pixels; right: Spatial mapping of UMAP clusters. 8 clusters were observed.

**Figure S4.**
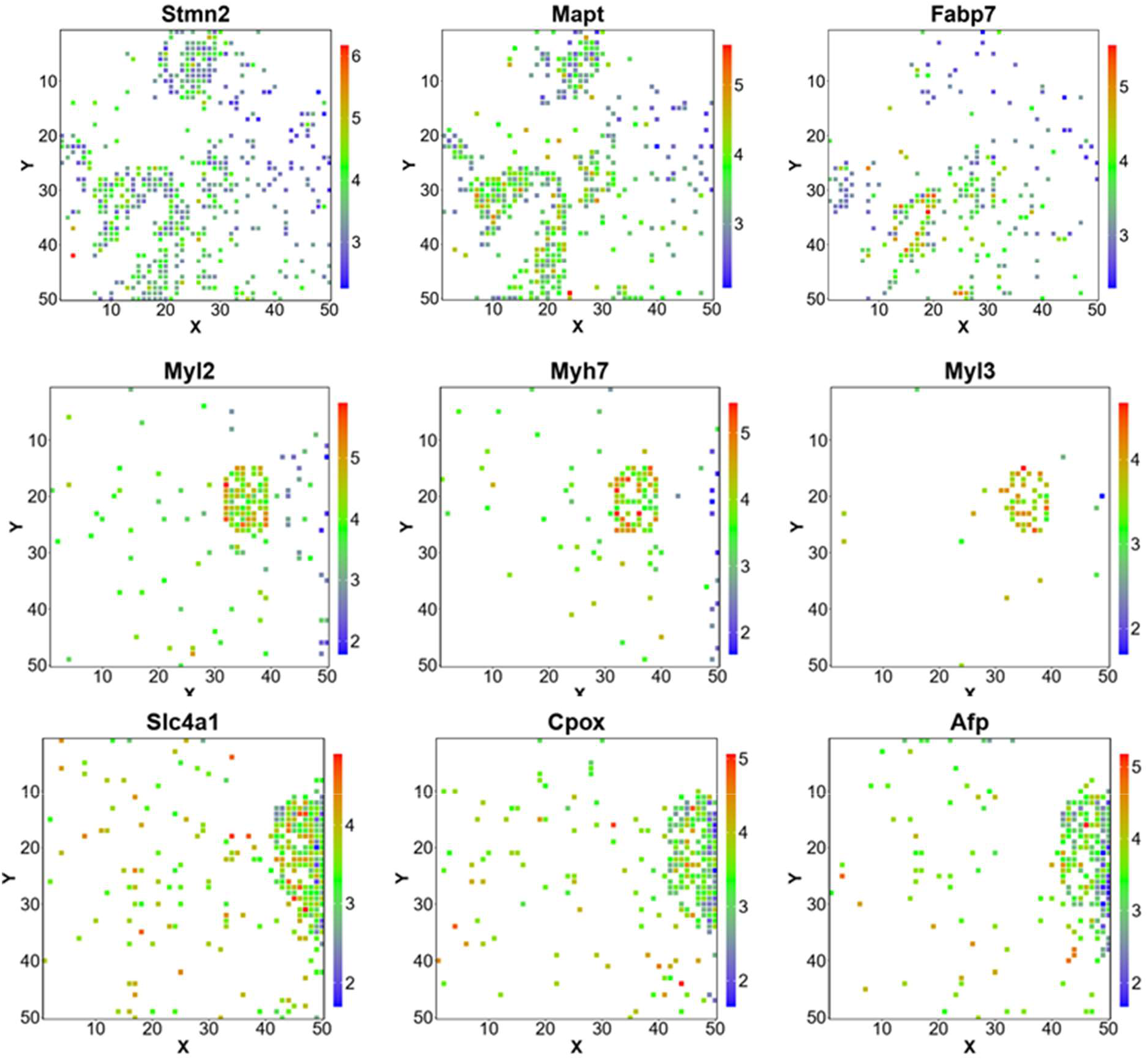
Spatial expression heatmap of select individual genes in FFPE-1 and FFPE-2.

**Figure S5.**
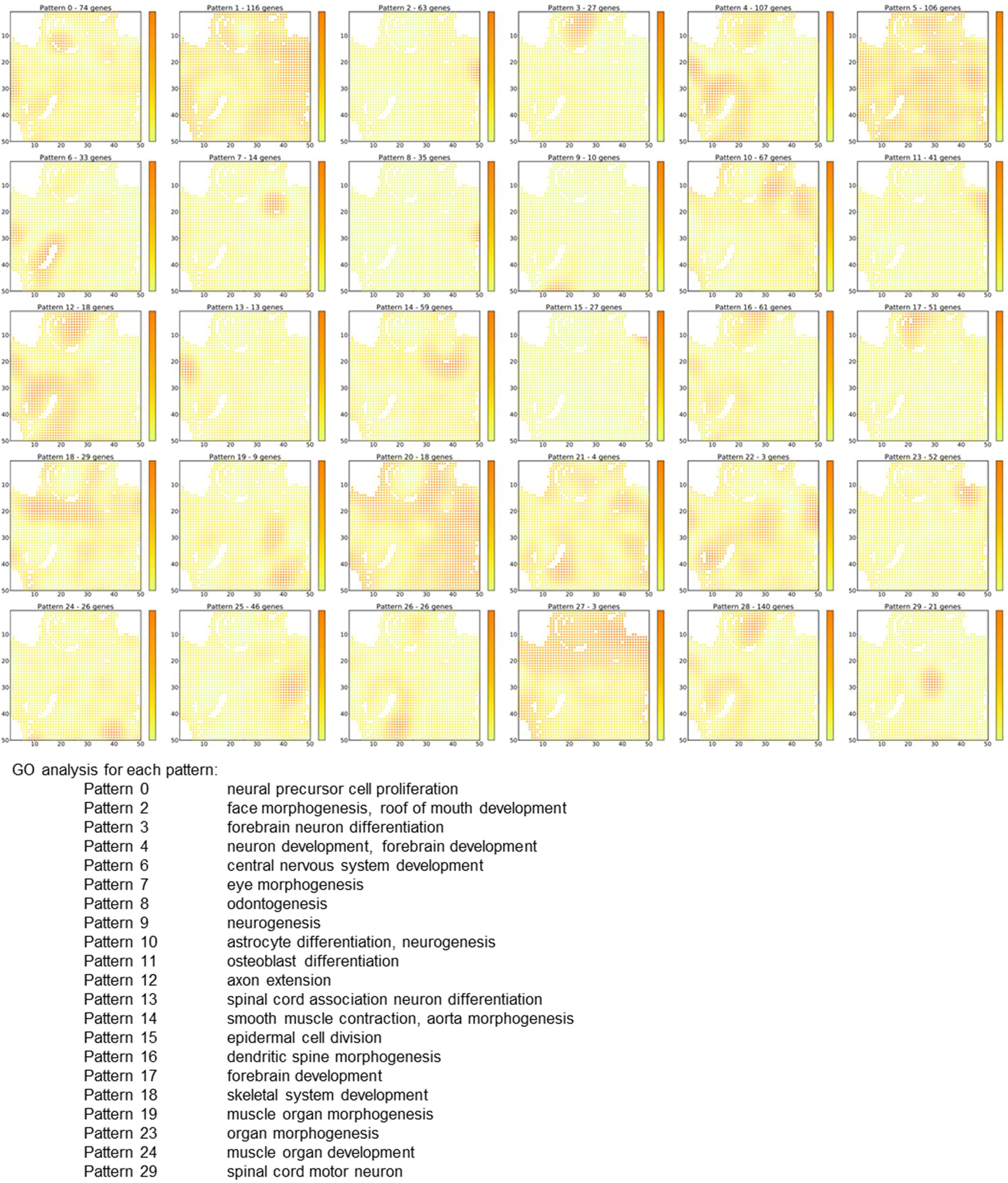
SpatialDE patterns of FFPE-1 tissue section. Spatial maps of 30 clusters were identified. Select clusters with distinct GO pathways are listed.

**Figure S6.**
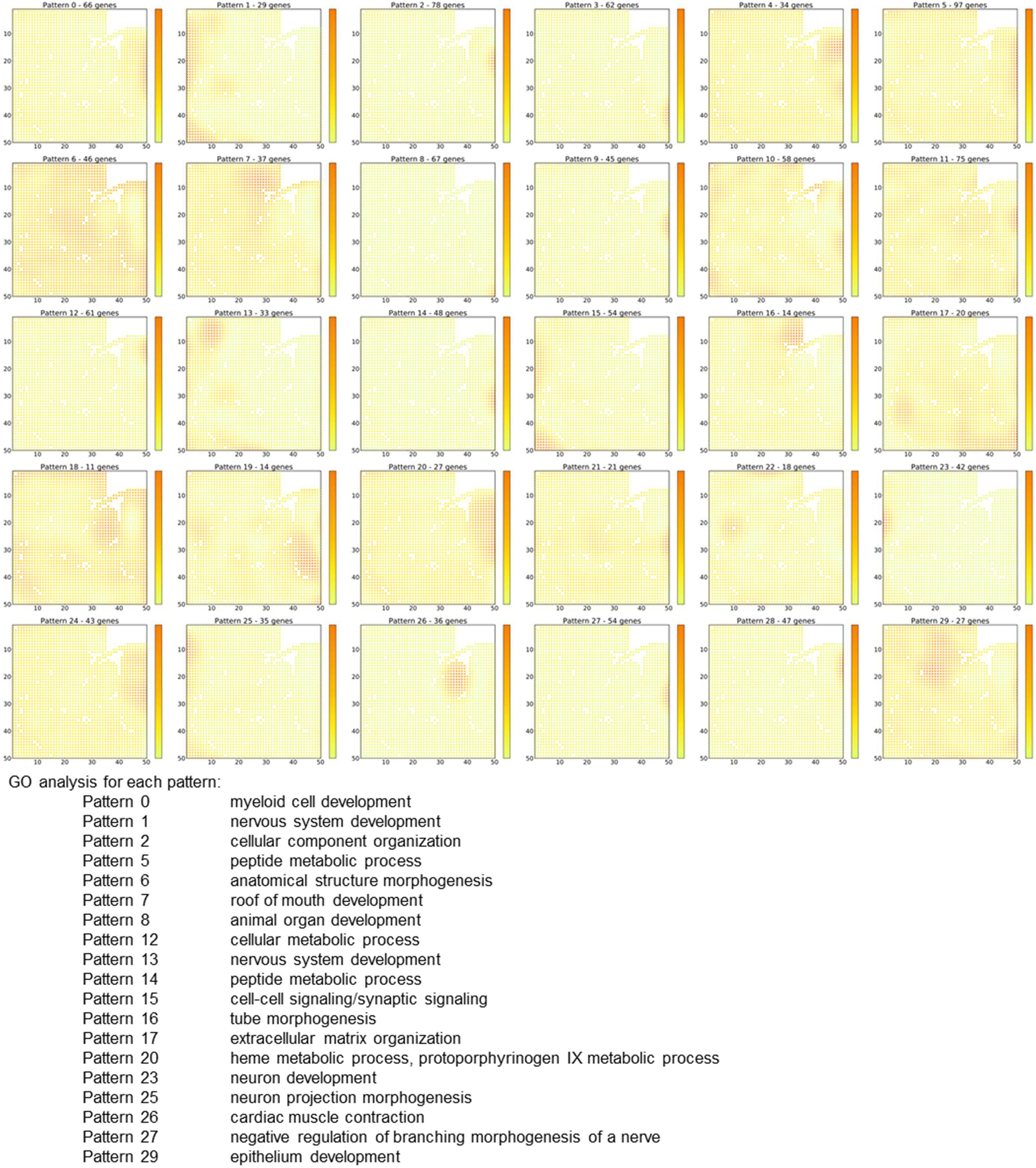
SpatialDE patterns of FFPE-2 tissue section. Spatial maps of 30 clusters were identified. Select clusters with distinct GO pathways are listed.

**Figure S7.**
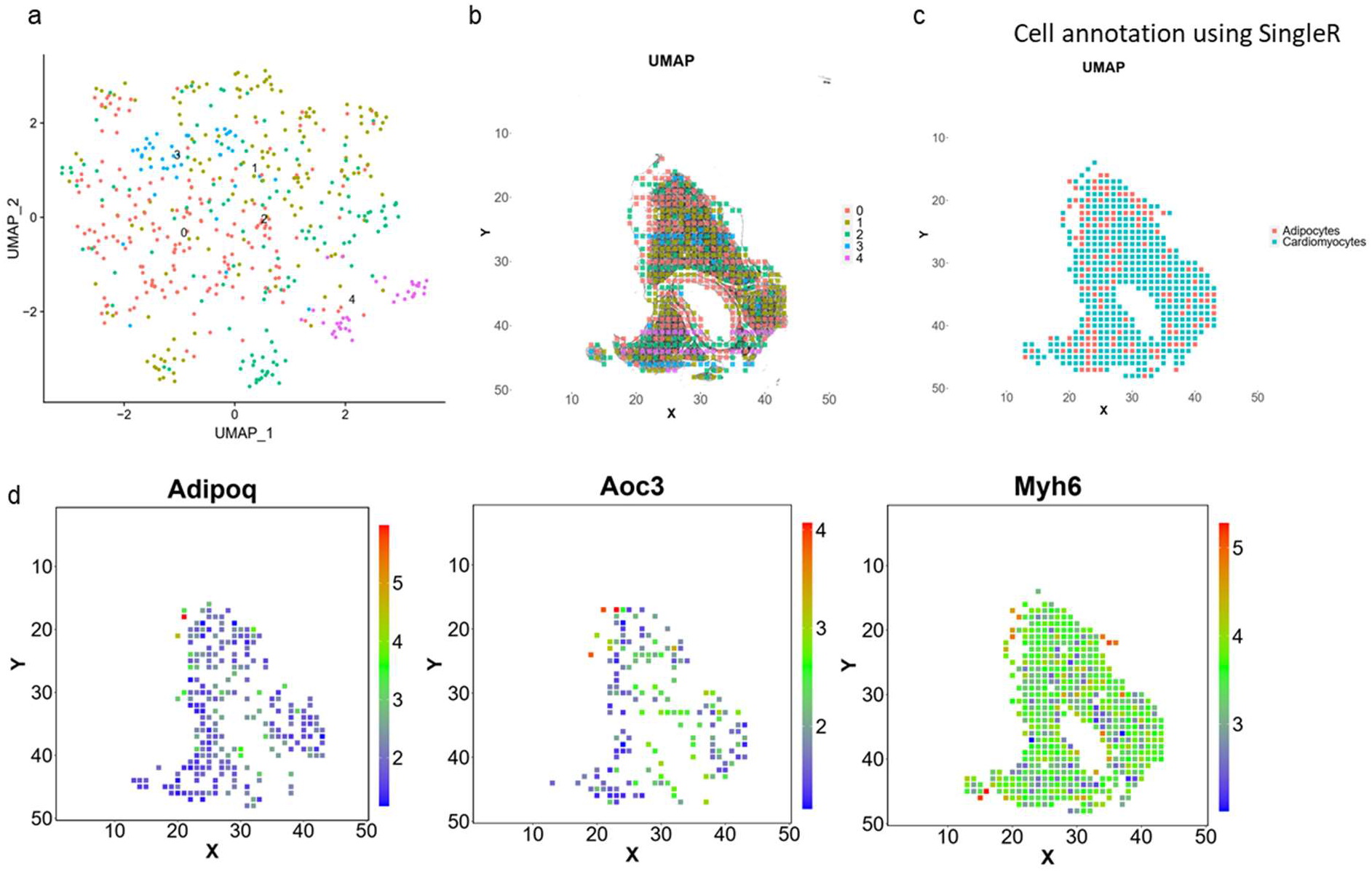
Clustering of the mouse aorta sample and visualization of individual genes. (a) UMAP clustering of all DBiT-seq pixels. (b) Spatial mapping of UMAP clusters in (a). (c) SingleR prediction of cell types using the built-in reference. (d) Spatial Mapping of individual genes: *Adipoq, Aoc3* and *Myh6*.

**Figure S8.**
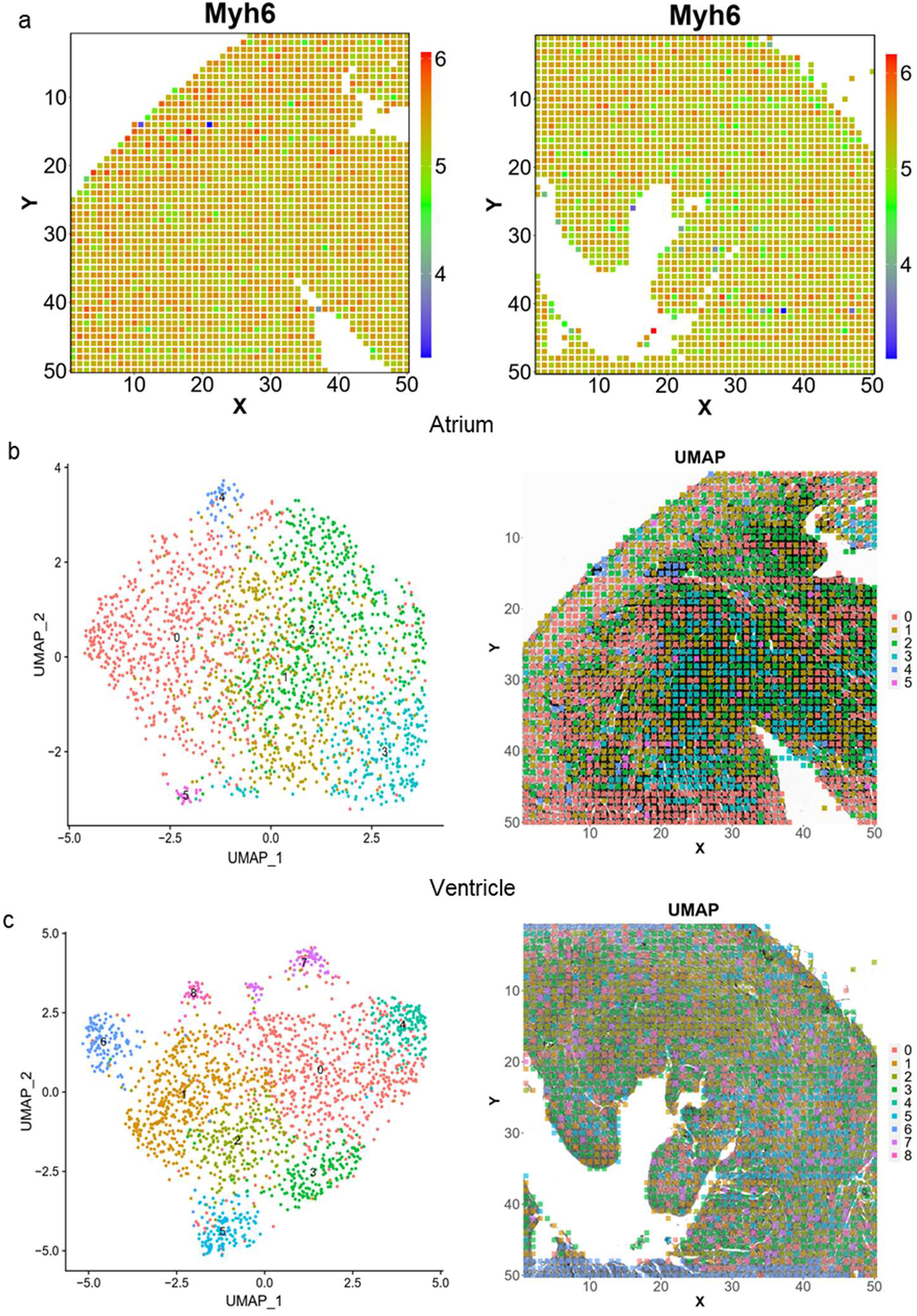
*My6* gene distribution and UMAP of atrium and ventricle. (a) *My6* gene in atrium (left) and ventricle (right). (b) Unsupervised clustering of atrium. (c) Unsupervised clustering of ventricle.

**Figure S9.**
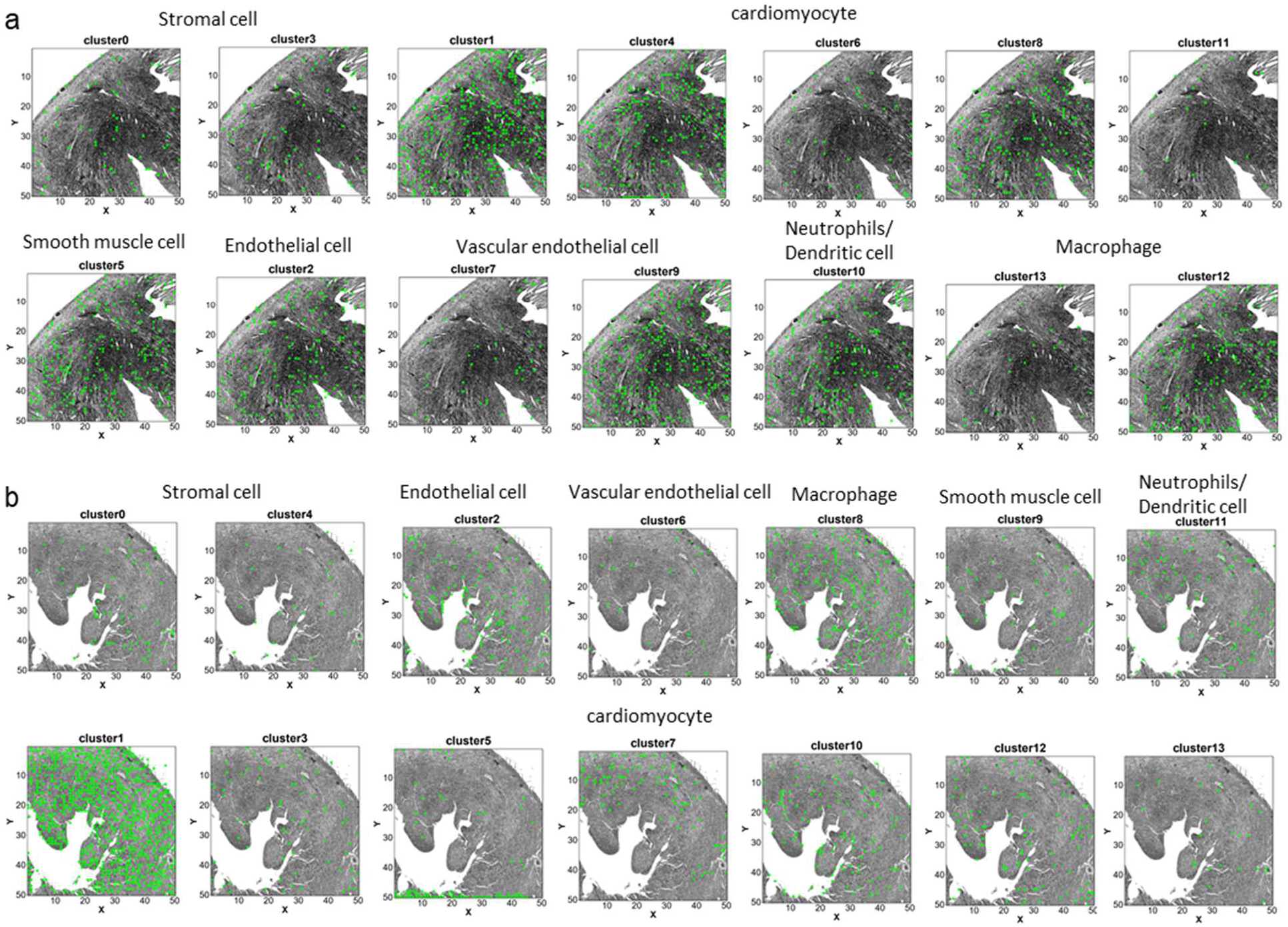
Spatial distribution of all 14 clusters in mouse atrium and ventricle. (a) cell distribution in mouse atrium. (b) cell distribution in mouse ventricle.

### Supplementary Tables

**Table S1.**
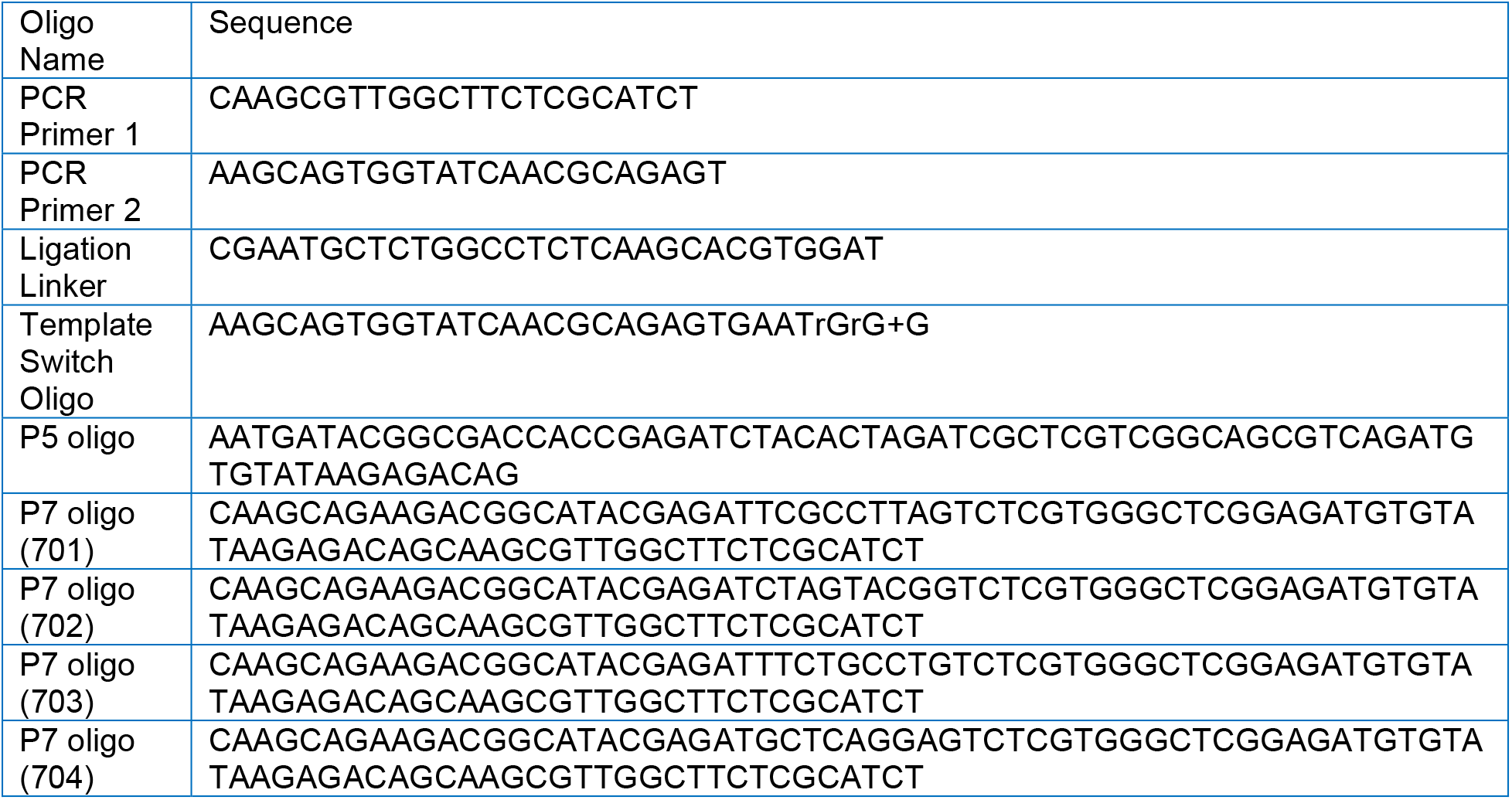
DNA oligos used for PCR and preparation of sequencing library.

**Table S2.**
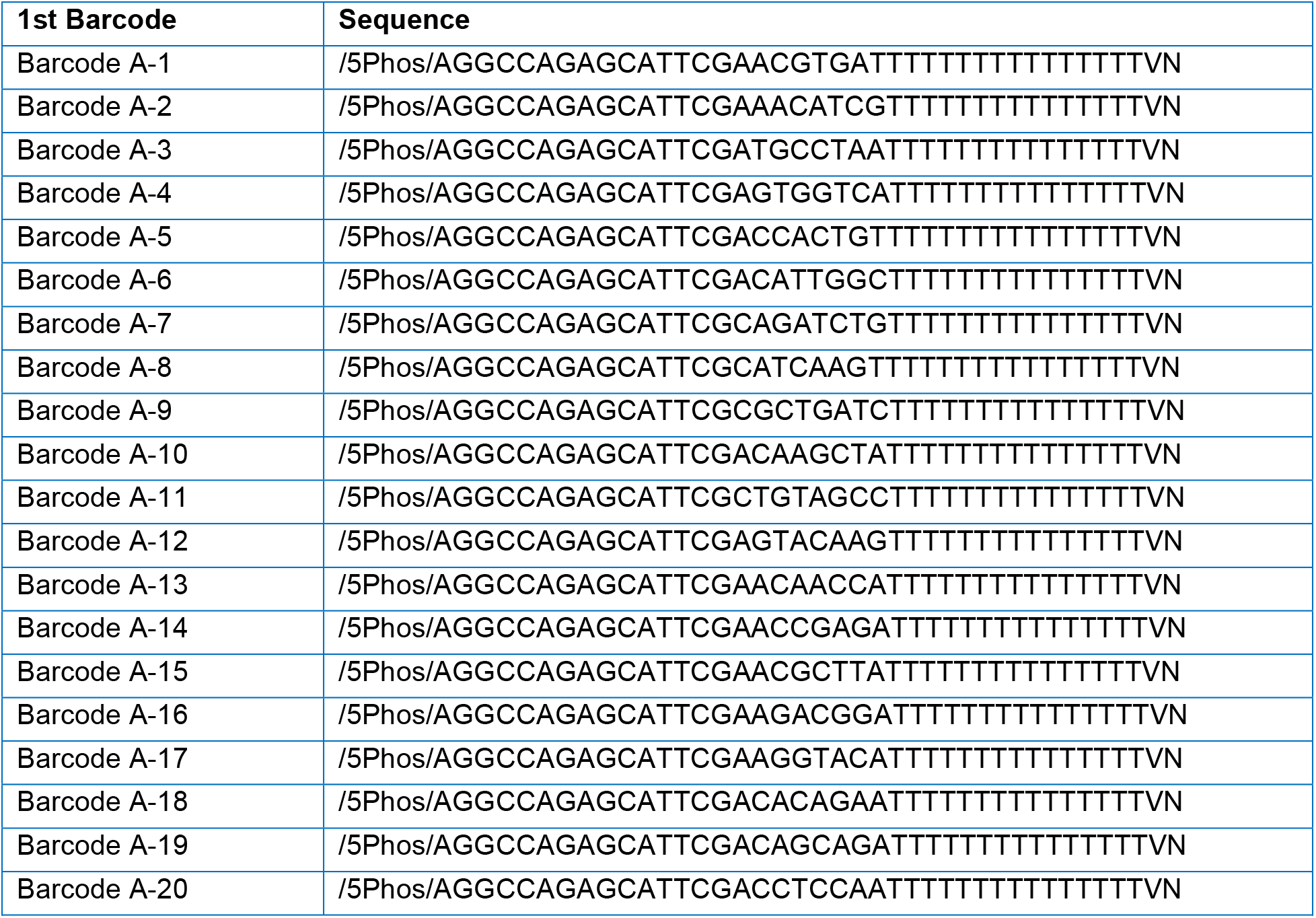
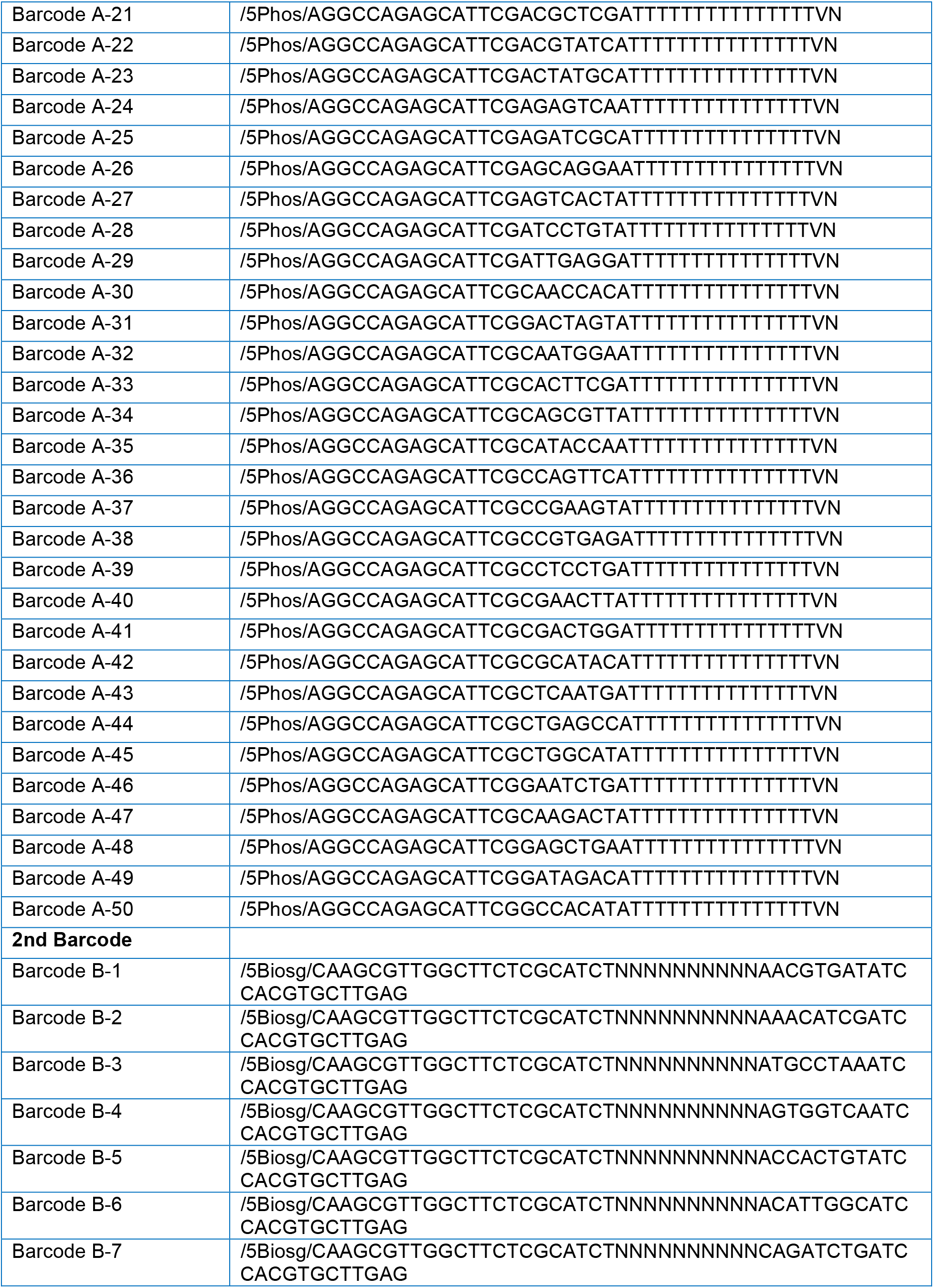
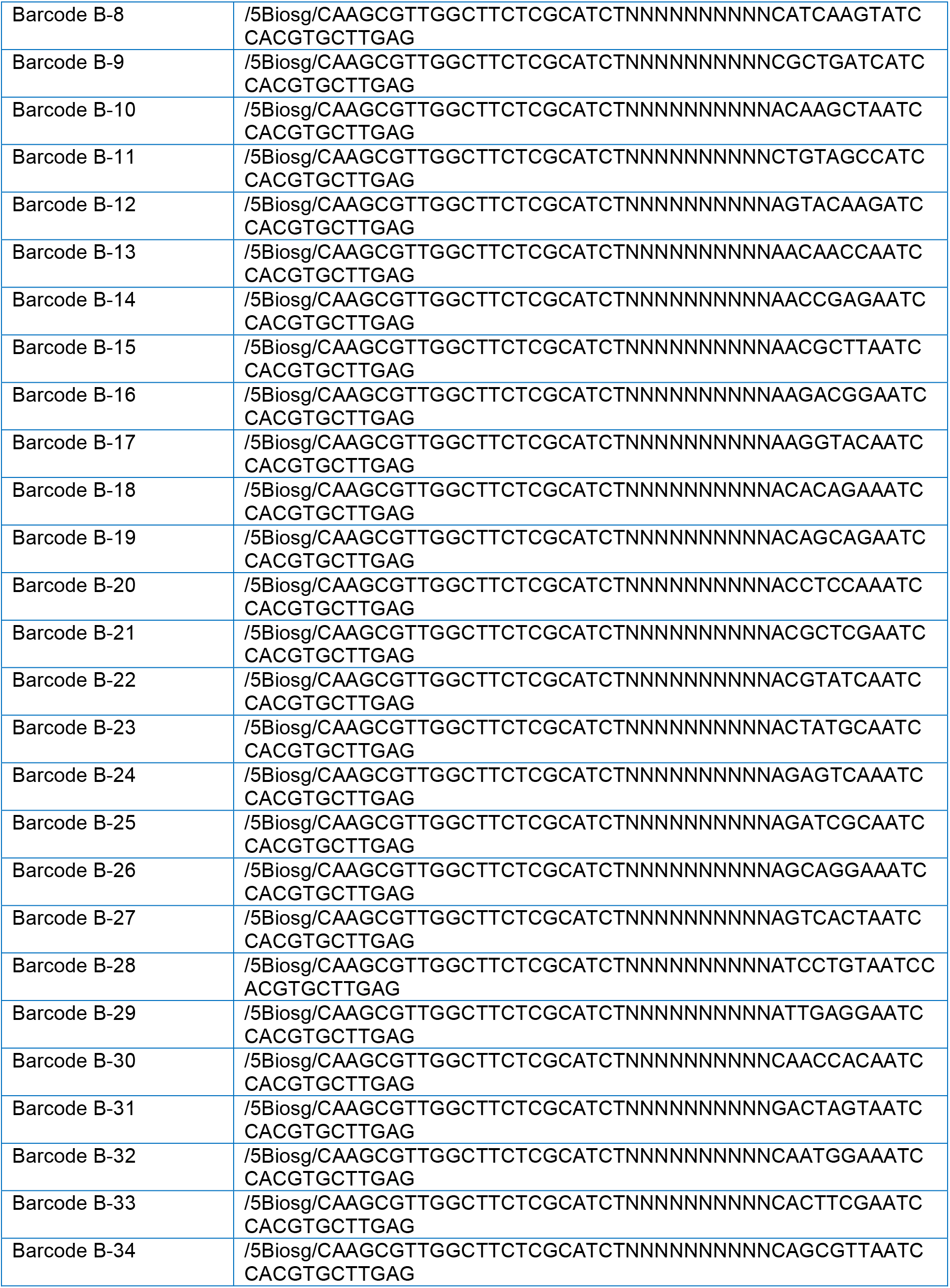
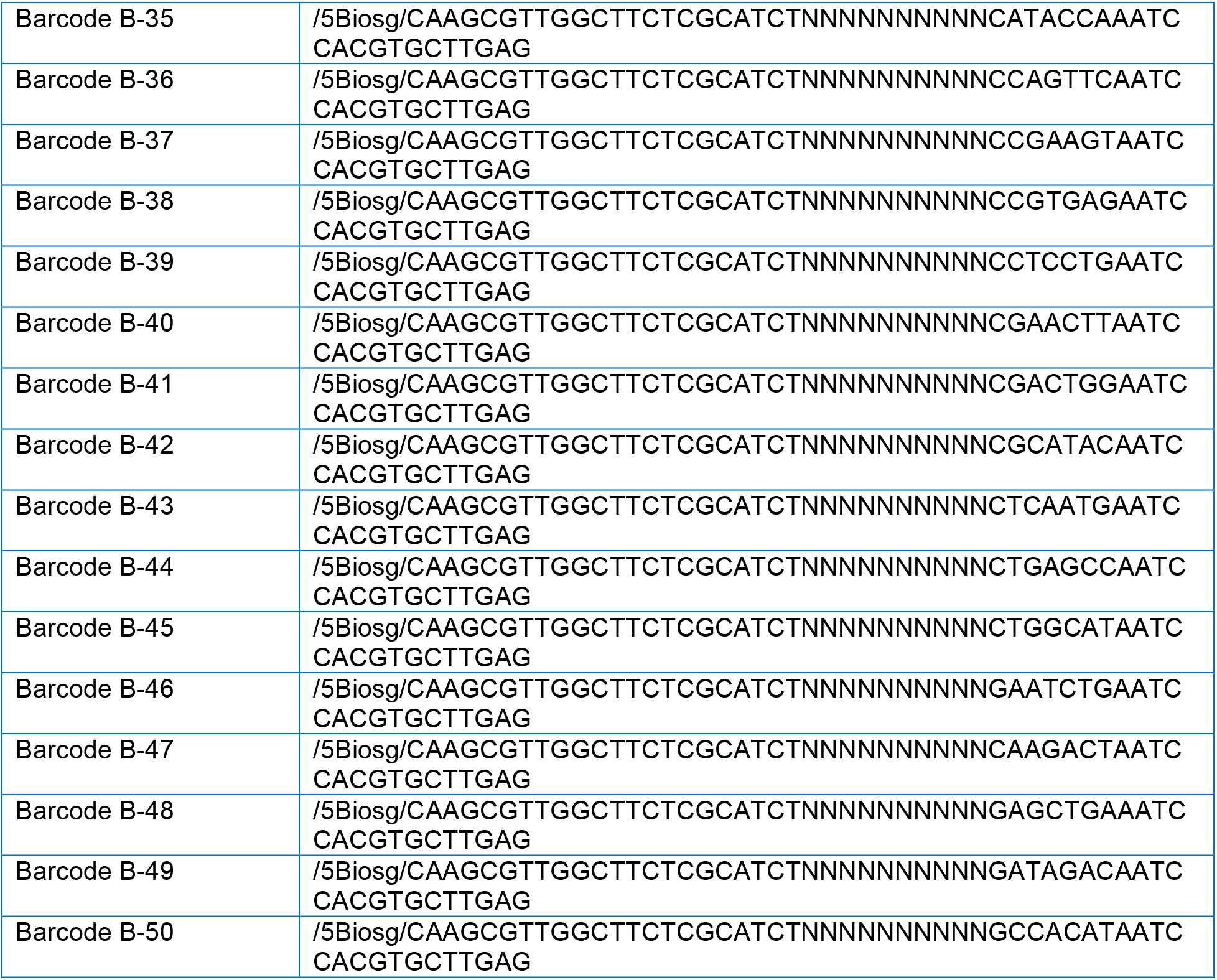
DNA barcode sequences.

**Table S3.** Chemicals and reagents.

| Reagent List |  |  |
| --- | --- | --- |
| Name | Cat. Number | Vender |
| Maxima H Minus | EP7051 | Thermo Fisher |
| dNTP mix | R0192 | Thermo Fisher |
| RNase Inhibitor | Y9240L | Enzymatics |
| SUPERase• In™ RNase Inhibitor | AM2694 | Thermo Fisher |
| T4 DNA Ligase | M0202L | New England Biolabs |
| Ampure XP beads | A63880 | Beckman Coulter |
| Dynabeads MyOne C1 | 65001 | Thermo Fisher |
|  | 65002 |  |
| Nextera XT DNA Preparation Kit | FC-131-1024 | Illumina |
| Kapa Hotstart HiFi ReadyMix | KK2601 | Kapa Biosystems |
| Proteinase K, recombinant, PCR grade | EO0491 | Thermo Fisher |
| **Buffer and chemical supplies** |  |  |
| RNAse free water | 10977015 | Invitrogen |
| Xylene | 534056-4L | Sigma |
| Ethanol | 187380-4L | Sigma |
| Formaldehyde solution | F8775-25ML | Sigma |
| Triton X-100 | T8787-100ML | Sigma |
| NEBuffer 3.1 | B7203S | New England Biolabs |
| T4 DNA Ligase Reaction Buffer | B0202S | New England Biolabs |
| Binding and washing (B&W) Buffer (2x) | 15568025 | Thermo Fisher |
|  | AM9261 | Thermo Fisher |
|  | AM9760G | Thermo Fisher |
| PMSF | 10837091001 | Sigma |
| Tween 20 | 3005 | Thermo Fisher |

## Methods section reference

Aran, D., Looney, A.P., Liu, L., Wu, E., Fong, V., Hsu, A., Chak, S., Naikawadi, R.P., Wolters, P.J., Abate, A.R., et al. (2019). Reference-based analysis of lung single-cell sequencing reveals a transitional profibrotic macrophage. Nat Immunol 20, 163–172.

Butler, A., Hoffman, P., Smibert, P., Papalexi, E., and Satija, R. (2018). Integrating single-cell transcriptomic data across different conditions, technologies, and species. Nat Biotechnol 36, 411–420.

Cao, J., Spielmann, M., Qiu, X., Huang, X., Ibrahim, D.M., Hill, A.J., Zhang, F., Mundlos, S., Christiansen, L., Steemers, F.J., et al. (2019). The single-cell transcriptional landscape of mammalian organogenesis. Nature 566, 496–502.

Navarro, J.F., Sjöstrand, J., Salmén F., Lundeberg, J., and Ståhl, P.L. (2017). ST Pipeline: an automated pipeline for spatial mapping of unique transcripts. Bioinformatics 33, 2591–2593.

Stuart, T., Butler, A., Hoffman, P., Hafemeister, C., Papalexi, E., Mauck III, W.M., Hao, Y., Stoeckius, M., Smibert, P., and Satija, R. (2019). Comprehensive integration of single-cell data. Cell 177, 1888–1902. e1821.

## References

1. Krafft, A.E., Duncan, B.W., Bijwaard, K.E., Taubenberger, J.K. & Lichy, J.H. Optimization of the Isolation and Amplification of RNA From Formalin-fixed, Paraffin-embedded Tissue: The Armed Forces Institute of Pathology Experience and Literature Review. Mol Diagn 2, 217–230 (1997).

2. Bass, B.P., Engel, K.B., Greytak, S.R. & Moore, H.M. A review of preanalytical factors affecting molecular, protein, and morphological analysis of formalin-fixed, paraffin-embedded (FFPE) tissue: how well do you know your FFPE specimen? Arch Pathol Lab Med 138, 1520–1530 (2014).

3. Hedegaard, J., Thorsen, K., Lund, M.K., Hein, A.-M.K., Hamilton-Dutoit, S.J., Vang, S., Nordentoft, I., Birkenkamp-Demtröder, K., Kruhøffer, M., Hager, H., Knudsen, B., Andersen, C.L., Sørensen, K.D., Pedersen, J.S., Ørntoft, T.F. & Dyrskjøt, L. Next-Generation Sequencing of RNA and DNA Isolated from Paired Fresh-Frozen and Formalin-Fixed Paraffin-Embedded Samples of Human Cancer and Normal Tissue. PLoS ONE 9, e98187 (2014).

4. Xie, R., Chung, J.-Y., Ylaya, K., Williams, R.L., Guerrero, N., Nakatsuka, N., Badie, C. & Hewitt, S.M. Factors Influencing the Degradation of Archival Formalin-Fixed Paraffin-Embedded Tissue Sections. J. Histochem. Cytochem. 59, 356–365 (2011).

5. Graw, S., Meier, R., Minn, K., Bloomer, C., Godwin, A.K., Fridley, B., Vlad, A., Beyerlein, P. & Chien, J. Robust gene expression and mutation analyses of RNA-sequencing of formalin-fixed diagnostic tumor samples. Sci Rep 5, 12335 (2015).

6. Sinicropi, D., Qu, K., Collin, F., Crager, M., Liu, M.-L., Pelham, R.J., Pho, M., Rossi, A.D., Jeong, J., Scott, A., Ambannavar, R., Zheng, C., Mena, R., Esteban, J., Stephans, J., Morlan, J. & Baker, J. Whole Transcriptome RNA-Seq Analysis of Breast Cancer Recurrence Risk Using Formalin-Fixed Paraffin-Embedded Tumor Tissue. PLoS ONE 7, e40092 (2012).

7. Beck, A.H., Weng, Z., Witten, D.M., Zhu, S., Foley, J.W., Lacroute, P., Smith, C.L., Tibshirani, R., van de Rijn, M., Sidow, A. & West, R.B. 3’-End Sequencing for Expression Quantification (3SEQ) from Archival Tumor Samples. PLoS ONE 5, e8768 (2010).

8. Klein, A.M., Mazutis, L., Akartuna, I., Tallapragada, N., Veres, A., Li, V., Peshkin, L., Weitz, D.A. & Kirschner, M.W. Droplet barcoding for single-cell transcriptomics applied to embryonic stem cells. Cell 161, 1187–1201 (2015).

9. Macosko, E.Z., Basu, A., Satija, R., Nemesh, J., Shekhar, K., Goldman, M., Tirosh, I., Bialas, A.R., Kamitaki, N., Martersteck, E.M., Trombetta, J.J., Weitz, D.A., Sanes, J.R., Shalek, A.K., Regev, A. & McCarroll, S.A. Highly Parallel Genome-wide Expression Profiling of Individual Cells Using Nanoliter Droplets. Cell 161, 1202–1214 (2015).

10. Kulkarni, A., Anderson, A.G., Merullo, D.P. & Konopka, G. Beyond bulk: a review of single cell transcriptomics methodologies and applications. Curr Opin Biotechnol 58, 129–136 (2019).

11. Lee, J.H., Daugharthy, E.R., Scheiman, J., Kalhor, R., Yang, J.L., Ferrante, T.C., Terry, R., Jeanty, S.S., Li, C., Amamoto, R., Peters, D.T., Turczyk, B.M., Marblestone, A.H., Inverso, S.A., Bernard, A., Mali, P., Rios, X., Aach, J. & Church, G.M. Highly multiplexed subcellular RNA sequencing in situ. Science 343, 1360–1363 (2014).

12. Lubeck, E., Coskun, A.F., Zhiyentayev, T., Ahmad, M. & Cai, L. Single-cell in situ RNA profiling by sequential hybridization. Nat Methods 11, 360–361 (2014).

13. Chen, K.H., Boettiger, A.N., Moffitt, J.R., Wang, S.Y. & Zhuang, X.W. Spatially resolved, highly multiplexed RNA profiling in single cells. Science 348 (2015).

14. Stahl, P.L., Salmen, F., Vickovic, S. et al. Visualization and analysis of gene expression in tissue sections by spatial transcriptomics. Science 353, 78–82 (2016).

15. Rodriques, S.G., Stickels, R.R., Goeva, A., Martin, C.A., Murray, E., Vanderburg, C.R., Welch, J., Chen, L.M., Chen, F. & Macosko, E.Z. Slide-seq: A scalable technology for measuring genome-wide expression at high spatial resolution. Science 363, 1463–1467 (2019).

16. Vickovic, S., Eraslan, G., Salmen, F., Klughammer, J., Stenbeck, L., Schapiro, D., Aijo, T., Bonneau, R., Bergenstrahle, L., Navarro, J.F., Gould, J., Griffin, G.K., Borg, A., Ronaghi, M., Frisen, J., Lundeberg, J., Regev, A. & Stahl, P.L. High-definition spatial transcriptomics for in situ tissue profiling. Nat Methods (2019).

17. Liu, Y., Yang, M., Deng, Y., Su, G., Enninful, A., Guo, C.C., Tebaldi, T., Zhang, D., Kim, D., Bai, Z., Norris, E., Pan, A., Li, J., Xiao, Y., Halene, S. & Fan, R. High-Spatial-Resolution Multi-Omics Atlas Sequencing of Mouse Embryos via Deterministic Barcoding in Tissue. bioRxiv, 788992 (2020).

18. Stickels, R.R., Murray, E., Kumar, P., Li, J., Marshall, J.L., Di Bella, D., Arlotta, P., Macosko, E.Z. & Chen, F. Sensitive spatial genome wide expression profiling at cellular resolution. bioRxiv, 2020.2003.2012.989806 (2020).

19. Svensson, V., Teichmann, S.A. & Stegle, O. SpatialDE: identification of spatially variable genes. Nat. Meth. 15, 343–346 (2018).

20. Cao, J., Spielmann, M., Qiu, X., Huang, X., Ibrahim, D.M., Hill, A.J., Zhang, F., Mundlos, S., Christiansen, L., Steemers, F.J., Trapnell, C. & Shendure, J. The single-cell transcriptional landscape of mammalian organogenesis. Nature 566, 496–502 (2019).

21. Hafemeister, C. & Satija, R. Normalization and variance stabilization of single-cell RNA-seq data using regularized negative binomial regression. bioRxiv, 576827 (2019).

22. Baron, M.H., Isern, J. & Fraser, S.T. The embryonic origins of erythropoiesis in mammals. Blood 119, 4828–4837 (2012).

23. Kalluri, A.S., Vellarikkal, S.K., Edelman, E.R., Nguyen, L., Subramanian, A., Ellinor, P.T., Regev, A., Kathiresan, S. & Gupta, R.M. Single-Cell Analysis of the Normal Mouse Aorta Reveals Functionally Distinct Endothelial Cell Populations. Circulation 140, 147–163 (2019).

24. Lynn Ray, J., Leach, R., Herbert, J.-M. & Benson, M. Isolation of vascular smooth muscle cells from a single murine aorta. Methods in Cell Science 23, 185–188 (2001).

25. Zhou, P. & Pu, W.T. Recounting Cardiac Cellular Composition. Circ. Res. 118, 368–370 (2016).

26. Schaum, N., Karkanias, J., Neff, N.F. et al. Single-cell transcriptomics of 20 mouse organs creates a Tabula Muris. Nature 562, 367–372 (2018).

27. Navarro, J.F., Sjöstrand, J., Salmén, F., Lundeberg, J. & Ståhl, P.L. ST Pipeline: an automated pipeline for spatial mapping of unique transcripts. Bioinformatics 33, 2591–2593 (2017).

28. Stuart, T., Butler, A., Hoffman, P., Hafemeister, C., Papalexi, E., Mauck III, W.M., Hao, Y., Stoeckius, M., Smibert, P. & Satija, R. Comprehensive integration of single-cell data. Cell 177, 1888–1902. e1821 (2019).

29. Butler, A., Hoffman, P., Smibert, P., Papalexi, E. & Satija, R. Integrating single-cell transcriptomic data across different conditions, technologies, and species. Nat. Biotechnol. 36, 411–420 (2018).

30. Aran, D., Looney, A.P., Liu, L., Wu, E., Fong, V., Hsu, A., Chak, S., Naikawadi, R.P., Wolters, P.J., Abate, A.R., Butte, A.J. & Bhattacharya, M. Reference-based analysis of lung single-cell sequencing reveals a transitional profibrotic macrophage. Nat. Immunol. 20, 163–172 (2019).

